# Persistent increases of PKMζ in memory-activated neurons trace LTP maintenance during spatial long-term memory storage

**DOI:** 10.1101/2020.02.05.936146

**Authors:** Changchi Hsieh, Panayiotis Tsokas, Alejandro Grau-Perales, Edith Lesburguères, Joseph Bukai, Kunal Khanna, Joelle Chorny, Ain Chung, Claudia Jou, Nesha S. Burghardt, Christine A. Denny, Rafael E. Flores-Obando, Benjamin Rush Hartley, Laura Melissa Rodríguez Valencia, A. Iván Hernández, Peter J. Bergold, James E. Cottrell, Juan Marcos Alarcon, André Antonio Fenton, Todd Charlton Sacktor

## Abstract

PKMζ is an autonomously active PKC isoform crucial for the maintenance of synaptic long-term potentiation (LTP) and long-term memory. Unlike other protein kinases that are transiently stimulated by second messengers, PKMζ is persistently activated through sustained increases in kinase protein expression. Therefore, visualizing increases in PKMζ expression during long-term memory storage might reveal the sites of its persistent action and thus the location of memory-associated LTP maintenance in the brain. Using quantitative immunohistochemistry validated by the lack of staining in PKMζ-null mice, we examined the amount and distribution of PKMζ in subregions of the hippocampal formation of wild-type mice during LTP maintenance and spatial long-term memory storage. During LTP maintenance in hippocampal slices, PKMζ increases in the pyramidal cell body and stimulated dendritic layers of CA1 for at least 2 h. During spatial memory storage, PKMζ increases in CA1 pyramidal cells for at least 1 month, paralleling the persistence of the memory. The subset of CA1 pyramidal cells that are tagged by immediate early gene *Arc*-driven transcription of fluorescent proteins, whose expression increases during initial memory formation, also expresses the persistent increase of PKMζ during memory storage. In the memory-tagged cells, the increased PKMζ expression persists in dendritic compartments within *stratum radiatum* for 1 month, indicating the long-term storage of information in the CA3-to-CA1 pathway during remote spatial memory. We conclude that persistent increases in PKMζ trace the molecular mechanism of LTP maintenance and thus the sites of information storage within brain circuitry during long-term memory.

## Introduction

The persistent action of PKMζ is a leading candidate for a molecular mechanism of long-term memory storage (Sacktor & Fenton, 2018). PKMζ is a nervous system-specific, atypical PKC isoform with autonomous enzymatic activity (Sacktor *et al.*, 1993). The autonomous activity of PKMζ is due to its unusual structure that differs from other PKC isoforms. Full-length PKC isoforms consist of two domains — a catalytic domain and an autoinhibitory regulatory domain that suppresses the catalytic domain. These full-length isoforms are inactive until second messengers bind to the regulatory domain and induce a conformational change that releases the autoinhibition. The second messengers that activate full-length PKCs, such as Ca^2+^ or diacylglycerol, have short half-lives, resulting in transient increases in kinase activity.

PKMζ, in contrast, has only a PKCζ catalytic domain, and the lack of an autoinhibitory regulatory domain results in autonomous and thus persistent kinase activity once it is synthesized. PKMζ mRNA is transcribed from an internal promoter within the PKCζ/PKMζ gene that is active only in neural tissue (Hernandez *et al.*, 2003), and the mRNA is transported to the dendrites of neurons (Muslimov *et al.*, 2004). Under basal conditions PKMζ mRNA is translationally repressed (Hernandez *et al.*, 2003). High-frequency afferent synaptic activity that occurs during LTP induction or learning derepresses the PKMζ mRNA translation, triggering new synthesis of PKMζ protein (Osten *et al.*, 1996; Hernandez *et al.*, 2003; Tsokas *et al.*, 2016; Hsieh *et al.*, 2017). The newly synthesized PKMζ then translocates to postsynaptic sites, where the autonomous kinase action on AMPAR-channel trafficking potentiates transmission of synaptic pathways that had been strongly activated, but not synaptic pathways that had been inactive, in a process known as “synaptic tagging and capture” (Sajikumar *et al.*, 2005; Palida *et al.*, 2015).

Once increased, the steady-state amount of PKMζ can remain elevated. Increased PKMζ kinase activity is sufficient to maintain LTP, strongly suggesting that it also maintains long-term memory (Ling *et al.*, 2002; Pastalkova *et al.*, 2006; Tsokas *et al.*, 2016). Quantitative immunoblotting of microdissected CA1 regions from hippocampal slices shows that the increase of PKMζ in LTP maintenance persists for several hours and correlates with the degree of synaptic potentiation (Osten *et al.*, 1996; Tsokas *et al.*, 2016). Likewise, immunoblotting of dorsal hippocampus following spatial long-term memory storage reveals the increase of PKMζ persists from days to weeks and correlates with the degree and duration of memory retention (Hsieh *et al.*, 2017).

The close association of PKMζ levels with synaptic potentiation motivates the central hypothesis of this work, that visualizing distribution of PKMζ expression may reveal sites of the physiological maintenance mechanism of LTP within the dorsal hippocampus during spatial long-term memory storage. In the hippocampus, the CA3→CA1 pathway is preferentially engaged for memory-related information processing, whereas the temporoammonic entorhinal cortex layer 3 (EC3)→CA1 pathway is preferentially engaged for processing information related to current perceptions (Brun *et al.*, 2002; Lisman, 2005; Colgin *et al.*, 2009; Pavlowsky *et al.*, 2017; Choi *et al.*, 2018; Dvorak *et al.*, 2018). We therefore predict that persistent increased expression of PKMζ in the CA1 cells that are allocated for a place-avoidance memory will preferentially identify CA3→CA1 Schaffer collateral synapses at CA1 *stratum radiatum*, but not the EC3→CA1 synapses at *stratum lacunosum moleculare*. Using quantitative immunohistochemistry, we determine the location of increased PKMζ during LTP maintenance and long-term memory storage of active place avoidance, a conditioned behavior that requires increases in PKMζ activity in the hippocampus and depends upon the hippocampus from 1 day to 1 month (Pastalkova *et al.*, 2006; Tsokas *et al.*, 2016; Hsieh *et al.*, 2017).

## Materials and Methods

### Reagents

The ζ-specific rabbit anti-PKMζ C-2 polyclonal antiserum (1:10,000 for immunohistochemistry; 1:20,000 for immunoblots) was generated as previously described (Hernandez *et al.*, 2003). The PKCζ/PKMζ-specific peptide (TLPPFQPQITDDYGL) corresponds to an epitope in an isoform-specific region in the catalytic domain of PKCζ/PKMζ and was synthesized by Quality Control Biochemicals (Hopkinton, MA). The peptide was coupled to bovine serum albumin (Pierce), mixed with Titermax Gold (CytRx Corp., Norcross, GA), and injected intramuscularly into female New Zealand rabbits. After 1–3 boosts at 4-week intervals, the antisera were affinity-purified on peptide-conjugated Sulfolink gel columns (Pierce). Actin (1:5000) mouse monoclonal Ab and other reagents unless specified were from Sigma. Protein concentrations were determined using the Bio-Rad RC-DC Protein Assay kit (Bio-Rad), with bovine serum albumin as standard.

### Animals

All procedures were performed in compliance with the Institutional Animal Care and Use Committees of the State University of New York, Downstate Health Sciences University, and New York University. All efforts were made to minimize animal suffering and to reduce the number of animals used.

### Hippocampal slice preparation and recording

Acute hippocampal slices (450 µm) from 2- to 4-month-old, male C57BL/6J mice were prepared with a McIlwain tissue slicer as previously described (Tsokas *et al.*, 2005; Tsokas *et al.*, 2019), and maintained in an Oslo-type interface recording chamber at 31.5 °C for at least 2 h before recording. A concentric bipolar stimulating electrode was placed in CA3a *stratum radiatum*, and field excitatory postsynaptic potentials (fEPSPs) were recorded with a glass extracellular recording electrode (2–5 MΩ) placed 400 µm from the stimulating electrodes in CA1 *stratum radiatum*. Synaptic efficiency was measured as the maximum slope of the fEPSP. High-frequency stimulation consisted of standard two 100 Hz 1-sec tetanic trains, at 25% of spike threshold, spaced 20 sec apart, which is optimized to produce a relatively rapid onset synthesis of PKMζ and protein synthesis-dependent late-phase LTP (Osten *et al.*, 1996; Tsokas *et al.*, 2005). The slope from 10-90% of the rise of the field excitatory postsynaptic potential (fEPSP) was analyzed on an IBM computer using the WinLTP data acquisition program (Anderson & Collingridge, 2007).

### Active place avoidance (APA)

Active place avoidance procedures were carried out as previously described (Tsokas *et al.*, 2016). A 3- to 6-month-old, male mouse was placed on a 40-cm diameter circular arena rotating at 1 rpm within a room. The mouse’s position was determined 30 times per second by video tracking from an overhead camera (Tracker, Bio-Signal Group). A transparent wall made from polyethylene terephtalate glycol-modified thermoplastic prevented the animal from jumping off the elevated arena surface. This task challenges the animal on the rotating arena to avoid an unmarked 60° sector shock zone that is defined by distal visual landmarks in the room. When the system detected the mouse in the shock zone for 500 ms, a mild constant current foot-shock (60 Hz, 500 ms, 0.2 mA), minimally sufficient to make the animal move, was delivered and repeated each 1500 ms until the mouse leaves the shock zone. During a training trial, the arena rotation periodically transported the animal into the shock zone, and the mouse was forced to actively avoid it. In addition, a pretraining trial and a posttraining memory retention test without shock were conducted for the time equivalent to a training session (unless otherwise specified). The time to first enter the shock zone in each trial was recorded as an index of between-session memory.

In the 1-day memory retention task (Fig. 2), the mouse first received a 30 min pretraining trial without shock, followed by three 30 min training trials with a 2 h inter-trial interval. The avoidance memory was tested 24 h later. In the 1-week memory retention task (Supplementary Fig. 3), the mouse first received a 30-min pretraining trial without shock on Day 1, followed by three daily 30-min training trials. Memory was tested 7 days after the last training trial. In the 1-month memory retention task (Figs. 3, 5, Supplementary Fig. 4), the mouse first received a 30-min pretraining trial without shock, followed by three 30-min training trials with 2-h intertrial interval. For the stronger 30-day memory (Figs. 3, Supplementary Fig. 4), another three training trials were conducted 10 days later, and memory was tested 30 days after the last training trial. For the minimally conditioned 30-day memory (Fig. 5), the mice only received the initial three training trials, and memory was tested 30 days after the last training trial. For all experiments, the control untrained mice had the equivalent exposure to the apparatus as trained mice but were not shocked.

**FIG. 1.**
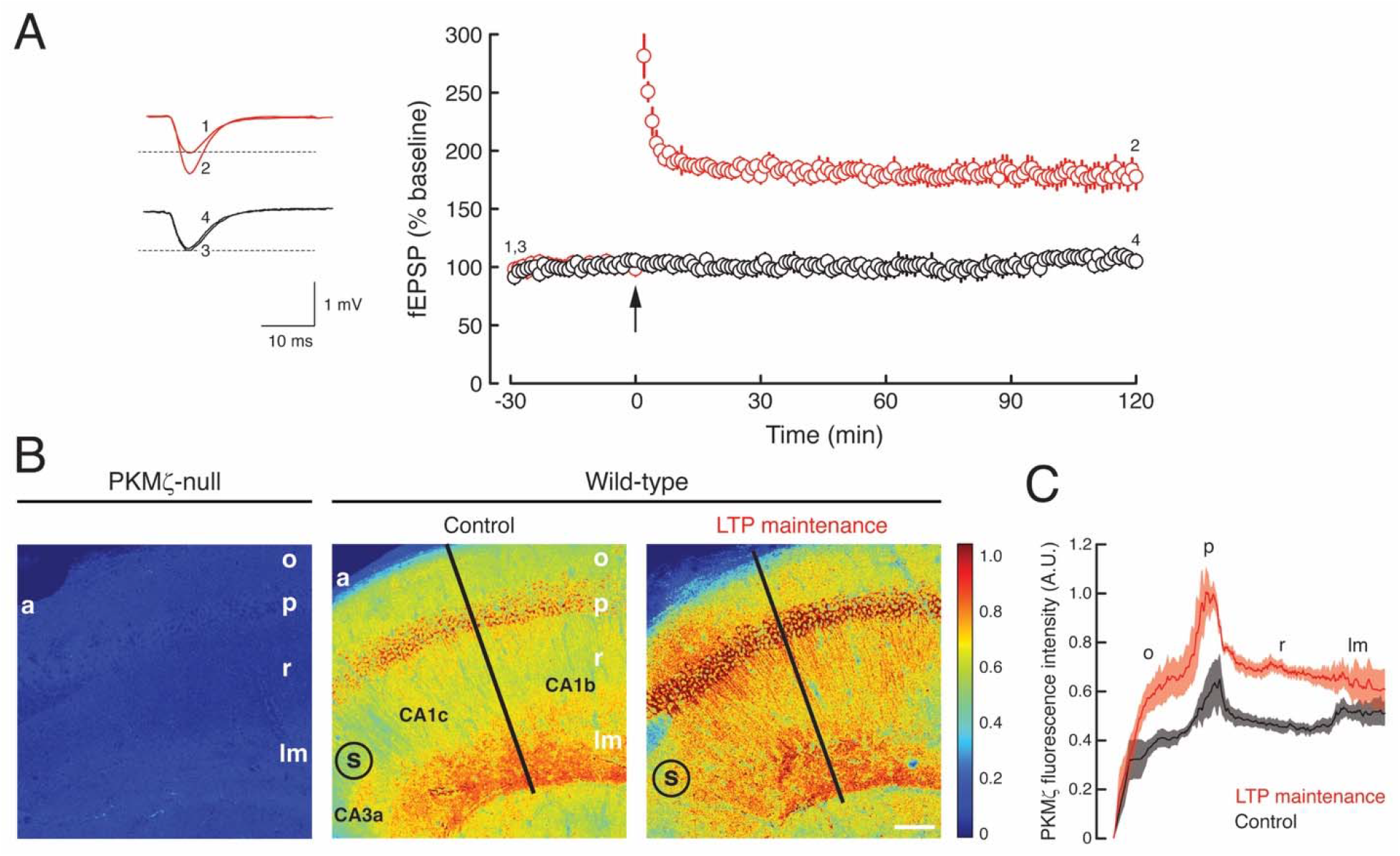
Persistent increased PKMζ in LTP maintenance. (A) Time-course of LTP (shown in red) and untetanized controls (shown in black). Left, representative fEPSPs correspond to numbered time points in time-course at right. Tetanization of Schaffer collateral/commissural afferent fibers is at arrow. Control slices are from the same hippocampus as tetanized slices and receive test stimulation for equivalent periods of time. (B) Representative immunohistochemistry of hippocampal CA1b/c subfields. Left, minimal background immunostaining in CA1 from PKMζ-null mouse. Middle, PKMζ in untetanized control slice from wild-type mouse, as recorded in (A). Right, persistent increased PKMζ 2 h post-tetanization in slice from wild-type mouse. Black lines correspond to location of profiles shown in (C). Colormap is shown as inset at right. Scale bar = 100 µm; s, placement of stimulating electrode at CA3/CA1 border; a, *alveus*; o, *stratum oriens*; p, *stratum pyramidale*; r, *stratum radiatum*; lm, *stratum lacunosum-moleculare*. (C) Profiles show increase in PKMζ in LTP maintenance (red), compared to untetanized controls (black). Mean ± SEM; n and statistics reported in Results. A.U., arbitrary units.

**FIG. 2.**
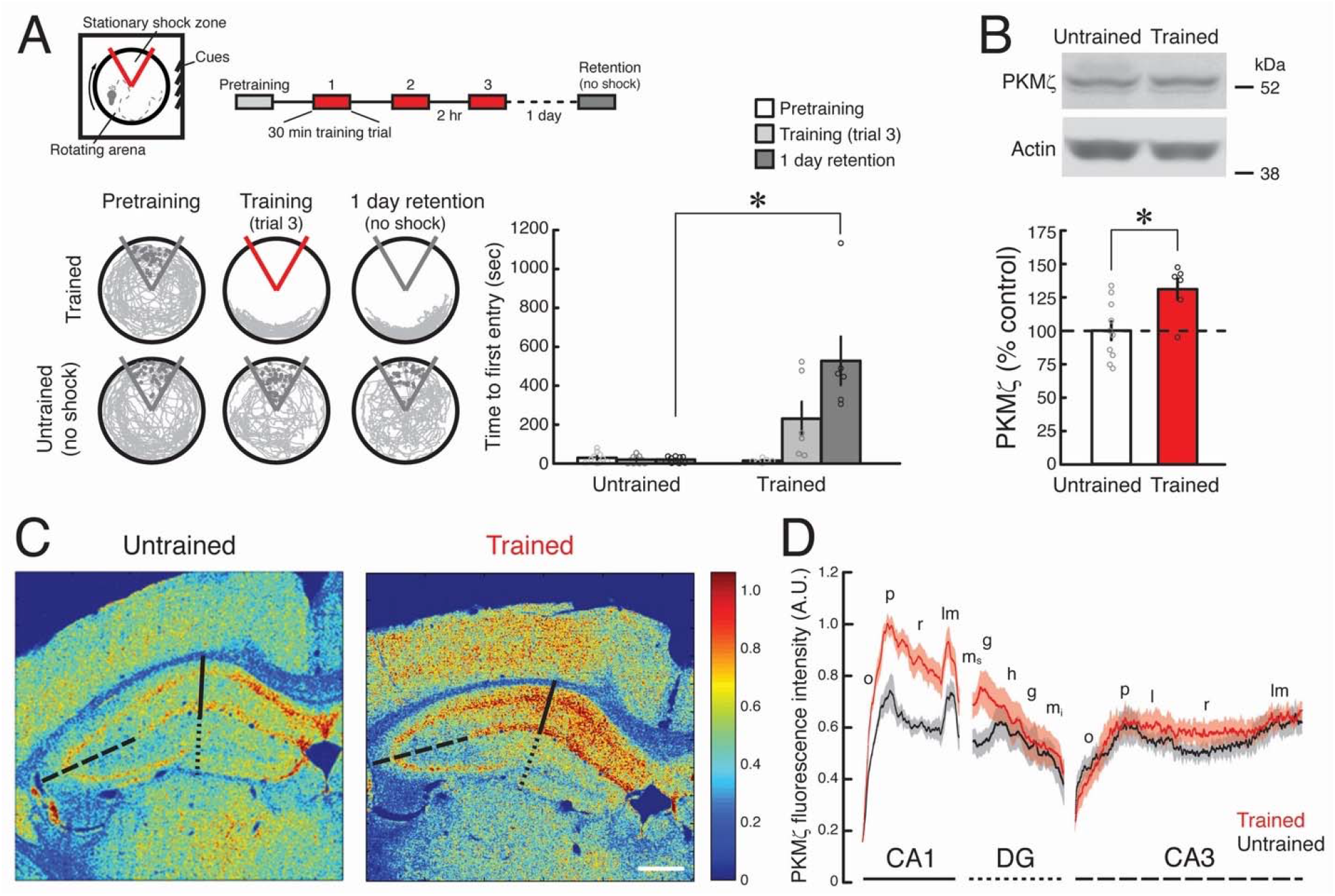
Persistent increased PKMζ in spatial long-term memory storage. (A) Above, left, schematic of active place avoidance apparatus. Above, right, schematic of spatial conditioning protocol. Active place avoidance conditioning produces spatial long-term memory at 1 day post-training, measured as increase in time to first entry into the shock zone. Below left, representative 10-min periods of paths at initiation of pretraining, at the end of training, and at initiation of retention testing with the shock off 1 day after training. The shock zone is shown in red with shock on, and gray with shock off. Gray circles denote where shocks would have been received if the shock were on. Right, time to first entry measure of active place avoidance memory (mean ± SEM of the data shown in circles; ^∗^, statistical significance, as reported in Results). (B) Immunoblots show PKMζ increases 1 day post-training in dorsal hippocampus. Above, representative immunoblots show increases in PKMζ between control untrained mouse and trained mouse, and actin as loading control. Below, mean data; the level of protein in the untrained control hippocampi is set to 100%. Mean ± SEM of the data shown in circles; ^∗^ denotes significance. (C) Representative immunohistochemistry shows PKMζ increases in dorsal hippocampus 1 day post-training. Left, image from control, untrained wild-type mouse shows widespread staining in hippocampal formation. Right, spatial training increases PKMζ immunostaining in CA1, but less in CA3 and DG. Black lines correspond to location of profiles shown in (D); CA1 (solid line), DG (dotted line), and CA3 (dashed line). Colormap is shown as insert at right. Scale bar = 500 µm. (D) Profiles show increases in PKMζ after training (red), compared to untrained controls (black), in CA1 and not in DG or CA3. S*trata*: o, *oriens*; p, *pyramidale*; r, *radiatum*; lm, *lacunosum-moleculare*; ms, *moleculare suprapyramidale*; g, *granule cell*; h, *hilus*; mi, *moleculare infrapyramidale*; l, *lucidum*. Mean data ± SEM; statistical significance reported in Results.

**FIG. 3.**
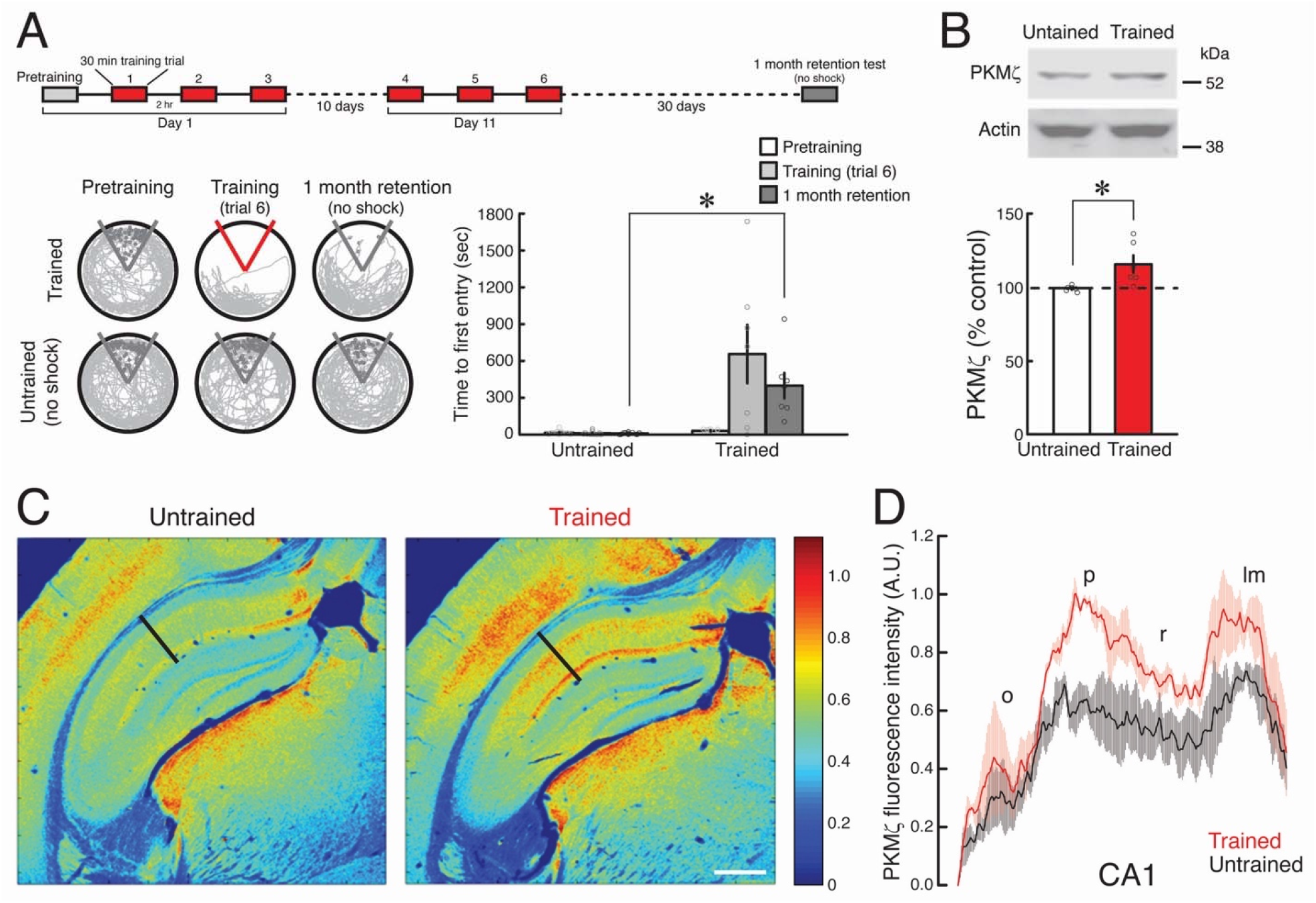
Persistent increased PKMζ in 1-month spatial remote memory storage. (A) Above, schematic of spatial conditioning protocol; repeated conditioning separated by 10 days produces remote memory lasting 30 days. Below left, representative 10-min periods of paths at initiation of pretraining, at the end of training, and at initiation of retention testing with the shock off 30 days after training. The shock zone is shown in red with shock on, and gray with shock off. Gray circles denote where shocks would have been received if the shock were on. Right, time to first entry measure of active place avoidance memory (mean ± SEM of the data shown in circles; ^∗^, statistical significance, as reported in Results). (B) Immunoblots show PKMζ increases 30 days post-training in dorsal hippocampus. Above, representative immunoblots show increases in PKMζ between control untrained mouse and trained mouse, and actin as loading control. Below, mean data with amount of PKMζ in the untrained control hippocampi set at 100%. Mean ± SEM of the data shown in circles; ^∗^ denotes significance. (C) Representative immunohistochemistry shows PKMζ in CA1 increases 30 days post-training. Black lines in untrained and trained mouse hippocampi correspond to locations of profiles shown in (D). Colormap is shown as insert at right. Scale bar = 500 µm. (D) Profiles show increase in PKMζ 30 days after training (red), compared to untrained controls (black). Mean data ± SEM; statistical significance reported in Results.

**FIG. 4.**
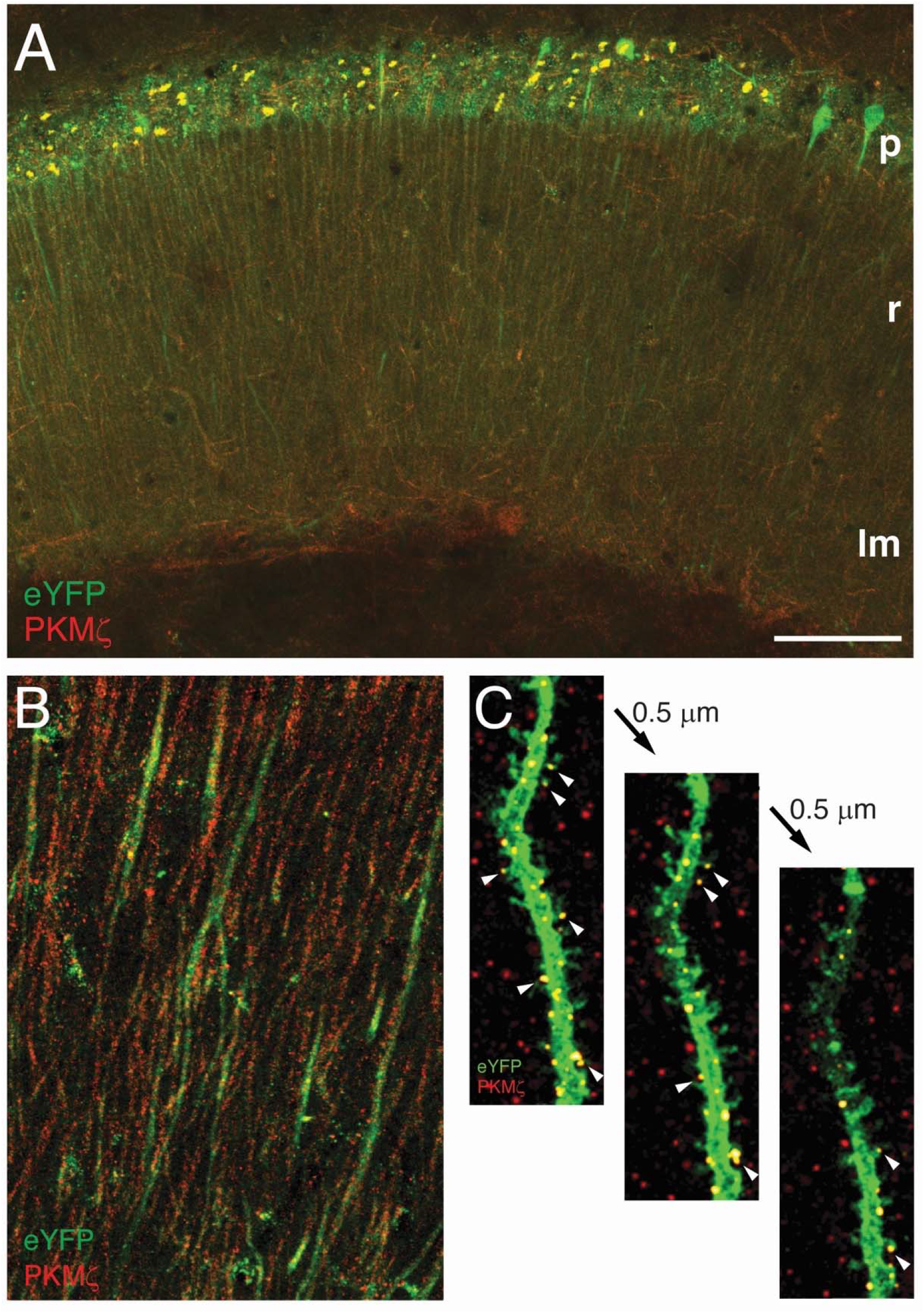
Colocalization of ChR2-eYFP and PKMζ in cell bodies and dendrites of memory-tagged CA1 pyramidal cells. (A) Representative image showing ChR2-eYFP (eYFP, green) expressed in response to *Arc* promoter-mediated Cre activation, PKMζ (red), and their overlap (yellow) in CA1; p, *strata pyramidale*; r, *radiatum*; lm, *lacunosum-moleculare*. (B) CA1 *stratum radiatum* shows PKMζ puncta both within eYFP+ dendrites of CA1 pyramidal cells, and outside of eYFP+ dendrites. Methods for A and B are as described for trained animals in experiments presented in Fig. 5. (C) PKMζ puncta in dendrite and spines (at arrowheads) of memory-tagged CA1 pyramidal cell. Three consecutive confocal images from a 0.5 µm Z-stack series of a CA1 dendrite from an *Arc*Cre-ChR2-eYFP mouse. The tissue was prepared for imaging by expansion microscopy as described in Methods. Scale bar = A: 100 µm, B: 30 µm, C: 6 µm.

**FIG. 5.**
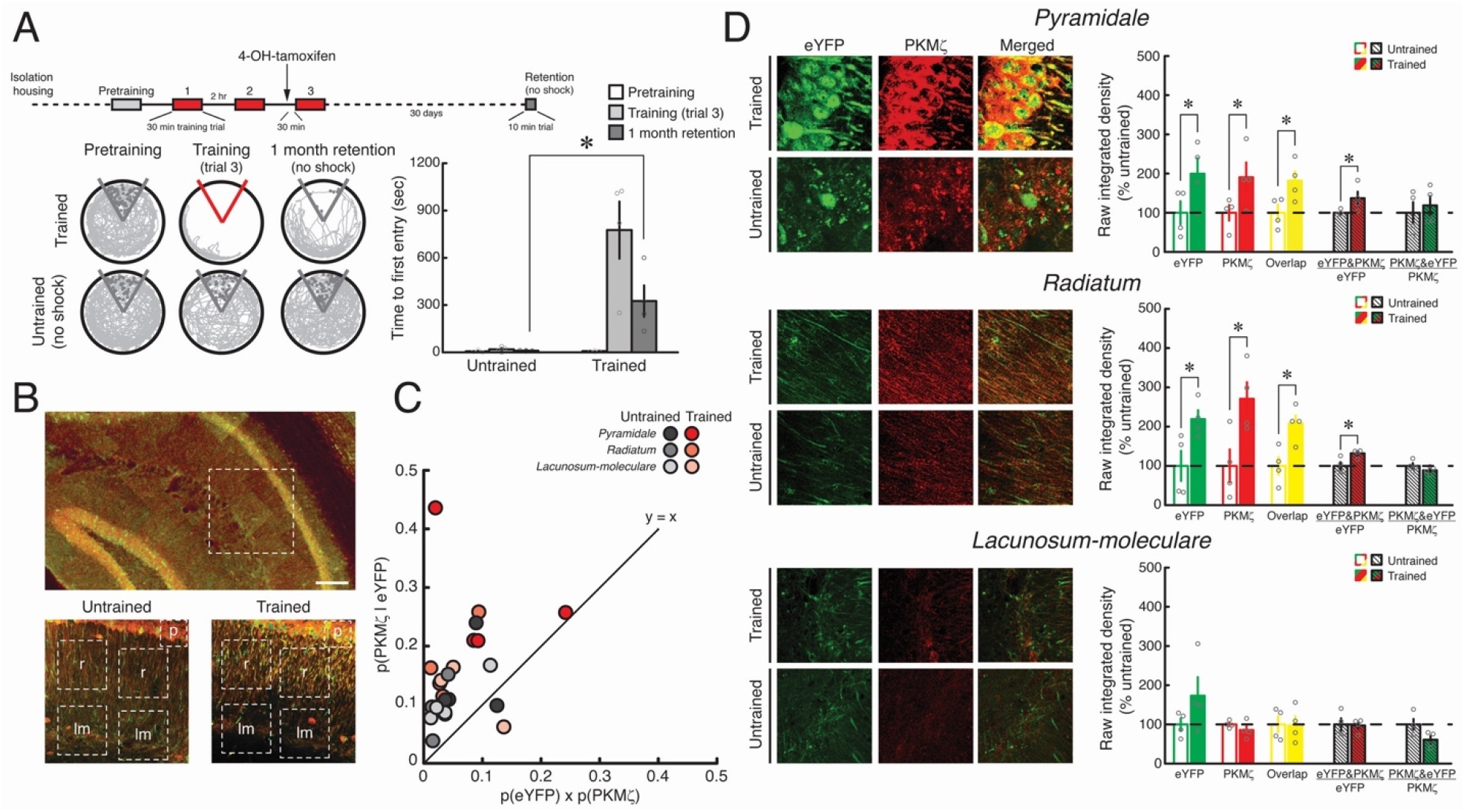
PKMζ increases in compartments of memory-tagged CA1 pyramidal cells persist for 1 month. (A) Above, schematic of spatial conditioning protocol producing memory lasting 30 days. Below left, representative 10-min periods of paths during initiation of pretraining, at the end of training, and at initiation of retention testing with the shock off 30 days after training. The shock zone is shown in red with shock on, and gray with shock off. Gray circles denote where shocks would have been received if the shock were on. Right, time to first entry measure of active place avoidance memory (mean ± SEM of the data shown in circles; ^∗^, statistical significance, as reported in Results). (B) Above, confocal image of hippocampus from trained *Arc*Cre-ChR2-eYFP mouse, showing colocalization (yellow) of eYFP (green) and PKMζ (red). White square shows representative region examined for analysis in images below. Below, white squares show representative regions of interest in r, *strata radiatum* and lm, *lacunosum-moleculare* analyzed in C and D. (C) Scatterplot shows the conditional probability of overlap p(PKMζ | eYFP) is greater than the product of p(PKMζ) and p(eYFP) for both trained and untrained mice in *strata pyramidale, radiatum*, and *lacunosum-moleculare*, supporting the hypothesis that experience increases PKMζ selectively in eYFP+ cells. (D) The intensities of the labeling for eYFP, PKMζ, and the overlapping signal are significantly increased in the trained group in *strata pyramidale* and *radiatum* for eYFP, for PKMζ, and for their overlap. No differences are found between the groups in *stratum lacunosum-moleculare*. Left, representative images; right, mean ± SEM of the data shown in circles with untrained control set at 100%; ^∗^ denotes significance. Manders coefficient M2 (eYFP&PKMζ / eYFP) estimates how much PKMζ is in the memory-tagged cells by computing the proportion of the eYFP signal that co-localizes with the PKMζ signal. M2 significantly increases in the trained group at *strata pyramidale* and *radiatum*, whereas no difference is observed at *stratum lacunosum-moleculare*. M1 (PKMζ&eYFP / PKMζ) estimates the proportion of the PKMζ signal that co-localizes with the eYFP signal. There is no group difference in M1 at any of the measured sites. Scale bar = B: upper panel, 150 µm; lower panels, 66 µm; D: *pyramidale*, 18 µm, *radiatum* and *lacunosum-moleculare*, 55 µm.

### Preparation of hippocampal extracts

Immediately after training, dorsal hippocampal extracts were prepared for immunoblotting. The hippocampi were removed after decapitation under deep isoflurane-induced anesthesia into ice-cold artificial cerebrospinal fluid with high Mg^2+^ (10 mM) and low Ca^2+^ (0.5 mM) (Sacktor *et al.*, 1993). The dorsal hippocampus consisting of half the hippocampus from the septal end was dissected out, snap-frozen, and stored in a microcentrifuge tube at −80°C until lysis. Dorsal hippocampi were lysed directly in the microcentrifuge tube using a motorized homogenizer, as previously described (Tsokas *et al.*, 2005; Tsokas *et al.*, 2016). Each dorsal hippocampus was homogenized in 100 µL of ice-cold modified RIPA buffer consisting of the following (in mM, unless indicated otherwise): 25 Tris-HCl (pH 7.4), 150 NaCl, 6 MgCl2, 2 EDTA, 1.25% NP-40, 0.125% sodium dodecyl sulfate (SDS), 0.625% Na deoxycholate, 4 *p*-nitrophenyl phosphate, 25 Na fluoride, 2 Na pyrophosphate, 20 dithiothreitol, 10 β-glycerophosphate, 1 µM okadaic acid, phosphatase inhibitor cocktail I & II (2% and 1%, respectively, Calbiochem), 1 phenylmethylsulfonyl fluoride, 20 µg/ml leupeptin, and 4 µg/ml aprotinin.

### Immunoblotting

The methods used were described previously (Tsokas *et al.*, 2005; Tsokas *et al.*, 2016). NuPage LDS Sample Buffer (4X) (Invitrogen, Carlsbad, CA) and β-mercaptoethanol were added to the homogenates, and the samples boiled for 5 min. The samples were then loaded (15-20 μg of protein per well) in a 4–20% Precast Protein Gel (Biorad) and resolved by SDS-polyacrylamide gel electrophoresis. Following transfer at 4 °C, nitrocellulose membranes (pore size, 0.2 μm; Invitrogen) were blocked for at least 30 min at room temperature with blocking buffer (5% non-fat dry milk in Tris-buffered saline containing 0.1% Tween 20 [TBS-T], or Licor Odyssey Blocking Buffer), then probed overnight at 4 °C using primary antibodies dissolved in blocking buffer or Licor Odyssey Blocking Buffer with 0.1% Tween 20 and 0.01% SDS. After washing in TBS-T (or phosphate-buffered saline + 0.1% Tween 20 [PBS-T]; 3 washes, 5 min each), the membranes were incubated with IRDye secondary antibodies (Licor). Proteins were visualized by the Licor Odyssey System. Densitometric analysis of the bands was performed using NIH ImageJ, and values were normalized to actin.

### Immunohistochemistry (IHC) and confocal microscopy

Brains were fixed by cardiac perfusion and hippocampal slices were placed in ice-cold 4% paraformaldehyde in 0.1 M phosphate buffer (PB, pH 7.4) immediately after behavioral testing or recording, and post-fixed for 48 h. Slices were then washed with PBS (pH 7.4) and cut into 40 µM sections using a Leica VT 1200S vibratome. As previously described (Tsokas *et al.*, 2005; Tsokas *et al.*, 2007), free-floating sections were permeabilized with PBS containing 0.3% Triton X-100 [PBS-TX100] for 1 h at room temperature and blocked with 10% normal donkey serum in PBS-TX100 (Blocking Buffer, BB) for 2.5 h at room temperature. The sections were then incubated overnight at 4 °C with primary antibody rabbit anti-PKMζ C-2 antisera (1:10,000) in BB. After washing 6 times for 10 mins each in PBS-TX100, the sections were then incubated with biotinylated donkey anti-rabbit secondary antibody (1:250 in BB; Jackson ImmunoResearch) for 2 h at room temperature. After washing 6 times for 10 min each in PBS-TX100, the sections were incubated with streptavidin conjugated-Alexa 647 (1:250 in PBS-TX100; Jackson ImmunoResearch) for 2 h at room temperature. After extensive washing with PBS-TX100, the sections were mounted with 4′,6-diamidino-2-phenylindole (DAPI) Fluoromount-G (Southern Biotech) or Vectashield (Vector Laboratories), and imaged using an Olympus Fluoview FV1000 or Zeiss LSM 800 confocal microscope at 4X, 10X, or 60X magnification. All parameters (pinhole, contrast and brightness) were held constant for all sections from the same experiment. To compare the intensity profile of PKMζ-immunostaining between images, we converted the images into grayscale and used the MATLAB functions *‘imread’* and *‘imagesc’* to scale the intensity distribution into the full range of a colormap. The same colormaps were used when comparing pairs of images, and representative colormaps are provided in the figures.

To compare brain sections between subjects, we normalized fluorescence intensity based upon the background level of staining in the absence of PKMζ in a PKMζ-null mouse. We compared the levels of fluorescence in the hippocampi of PKMζ-null mice and wild-type mice immunostained in parallel. The results show that: 1) the alveus has the lowest levels of immunostaining in wild-type mouse hippocampus, in line with previous work in rat hippocampus (Hernandez *et al.*, 2014), and 2) the low fluorescence signal is indistinguishable between the alveus of wild-type mice and PKMζ-null mice (Fig. 1B). Therefore, the background fluorescence level was set in the alveus of wild-type untetanized slices or untrained mice in all comparisons between tetanized and untetanized hippocampal slices and between pairs of trained and untrained mice.

The maximum intensity of immunostaining within the profile lines of pairs of trained and untrained mice or tetanized and untetanized slices was set to 1 (a representative set of images with profile lines is shown in Figs. 1-3, Supplementary Figs. 3, 4). The mean ± SEM of these sets of profile lines is presented in the figures as mean profile lines, with the maximum intensity of immunostaining of the averaged lines set to 1. The profile line in CA1 is set in the center of CA1b for behavioral experiments and at the border of CA1c and b for LTP experiments to visualize the projection fields of the tetanically stimulated afferents from CA3a (Amaral & Witter, 1989).

### Genetic tagging of memory-activated cells: ArcCreER^T2^ x ChR2-eYFP mice and ArcCreER^T2^ x eYFP mice

*Arc*CreER^T2^(+) (Denny *et al.*, 2014) x R26R-STOP-floxed-ChR2-eYFP (enhanced yellow fluorescent protein) (Srinivas *et al.*, 2001) and *Arc*CreER^T2^(+) x R26R-STOP-floxed-eYFP homozygous female mice were bred with R26R-STOP-floxed-ChR2-eYFP and R26R-STOP-floxed-eYFP (Perusini *et al.*, 2017; Lacagnina *et al.*, 2019) homozygous male mice, respectively (Srinivas *et al.*, 2001). All experimental mice were *Arc*CreER^T2^(+) and homozygous for the Chr2-eYFP or eYFP reporter. Genotyping was performed as previously described (Denny *et al.*, 2014).

### ArcCreER^T2^ x ChR2-eYFP mice: Behavior

Trained and untrained *Arc*-ChR2-eYFP mice were isolated in a customized, ventilated, temperature-, humidity-, and light-controlled (12/12 light/dark cycle), sound-attenuated chamber (Lafayette Instruments) for 3-4 days prior to pretraining. Precautions to prevent disturbances to the mice during the isolation housing were taken in order to reduce off-target labeling.

Pretraining for 30 min was followed 2 h later by three 30-min training trials with shock and a 2 h intertrial interval; untrained mice underwent the same protocol without shock. Following each trial, the mice were returned to the isolation chamber. Thirty minutes before the third trial when the task was familiar, and the trained mice demonstrated strong active avoidance memory, all mice (both trained and untrained) were injected intraperitoneally with 4-OH-tamoxifen (200-300 µl of 10 µg/µl, 0.01% of body weight) and returned to the home cage in the isolation chamber until the third trial. One month after training, the mice were returned to the arena and a 10-min trial was administered with no shock to evaluate memory retention. The mice were sacrificed after training and transcardially perfused with ice-cold 4% paraformaldehyde (PFA), and the brains were extracted and prepared for IHC.

### ArcCreER^T2^ x ChR2-eYFP mice: IHC

Thirty-µm slices of dorsal hippocampus were prepared from each animal, and IHC was performed for ChR2-eYFP (chicken anti-GFP antibody [1:1000], Abcam) and PKMζ, as described above. Three slices from each mouse were analyzed using ImageJ (version 1.53a; https://imagej.nih.gov/ij/; Rasband, W.S., ImageJ, U. S. National Institutes of Health, Bethesda, Maryland, USA, 1997-2018). For each slice, 8.5 µm-thick Z-stacks of the dorsal CA1 region were created using the ImageJ maximum intensity projection function. In each slice, two square regions of interest (ROI) centered on *strata pyramidale, radiatum, lacunosum-moleculare*, and *alveus* were examined, such that 6 measurements were made from each mouse in each ROI. Each image was normalized by the average value of PKMζ in the *alveus* to account for potential differences in processing, labeling intensity, and slice thickness across the mice. The raw integrated density (defined as the sum of the values for all pixels) of the ROI expressing the fluorescent label was measured for green (ChR2-eYFP) and red (PKMζ) pixels, and the average of the 6 (3 slices x 2 ROIs) normalized measurements was taken as representative for each mouse. We examined the cumulative distributions of the intensity of the pixels for the three channels (green, red, and yellow) in each area (data not shown), and by inspection there were no differences in the distributions between the trained and untrained animals in each brain area. The overlap between the ChR2-eYFP and PKMζ signals was calculated by measuring the raw integrated density of all the yellow pixels and dividing it by the total raw integrated density of the image (all green and red pixels combined). To assess labeling in trained mice relative to untrained mice, the representative values for each mouse were normalized by the corresponding average value for the untrained mice such that the average untrained value is 100%.

The percent area of the ROI that expresses the fluorescent label was also examined using the same threshold parameters as above with the average of the 6 measurements taken as representative for each mouse. To assess labeling in trained mice relative to untrained mice, the representative values for each mouse were normalized by the corresponding average value for the untrained mice such that the average untrained value is 100%. Manders coefficients corresponding to the proportion of ChR2-eYFP overlapping with PKMζ (M1) and the proportion of PKMζ overlapping with ChR2-eYFP (M2) were determined using the JACOP plugin for ImageJ. If M1 = 1, then eYFP expresses everywhere PKMζ expresses, and if M2 = 1, then PKMζ is expressed wherever eYFP is expressed. Since M1 and M2 vary according to the somato-dendritic compartment, to assess whether memory training changes the colocalization of eYFP and PKMζ at a specific compartment, we report the values for M1 and M2 relative to the corresponding average untrained control values.

### ArcCreER^T2^ x eYFP mice: Behavior

For labeling of principal cell soma (Fig. 6, Supplementary Fig. 5), mice were housed for 3 days prior to training in the isolation chambers to minimize off-target labeling of cells. Pretraining exposed mice to the behavioral room and arena, and was followed by 2 days of training (two 30-min trials per day with 40-min intertrial interval), followed 1 day later by a 10-min memory retention test without shock. At the end of each daily behavioral procedure, mice were returned to the isolation chamber. To label neurons active during memory retention, we injected the mice with 4-OH-tamoxifen to induce eYFP expression 30 min prior to the retention test session.

**FIG. 6.**
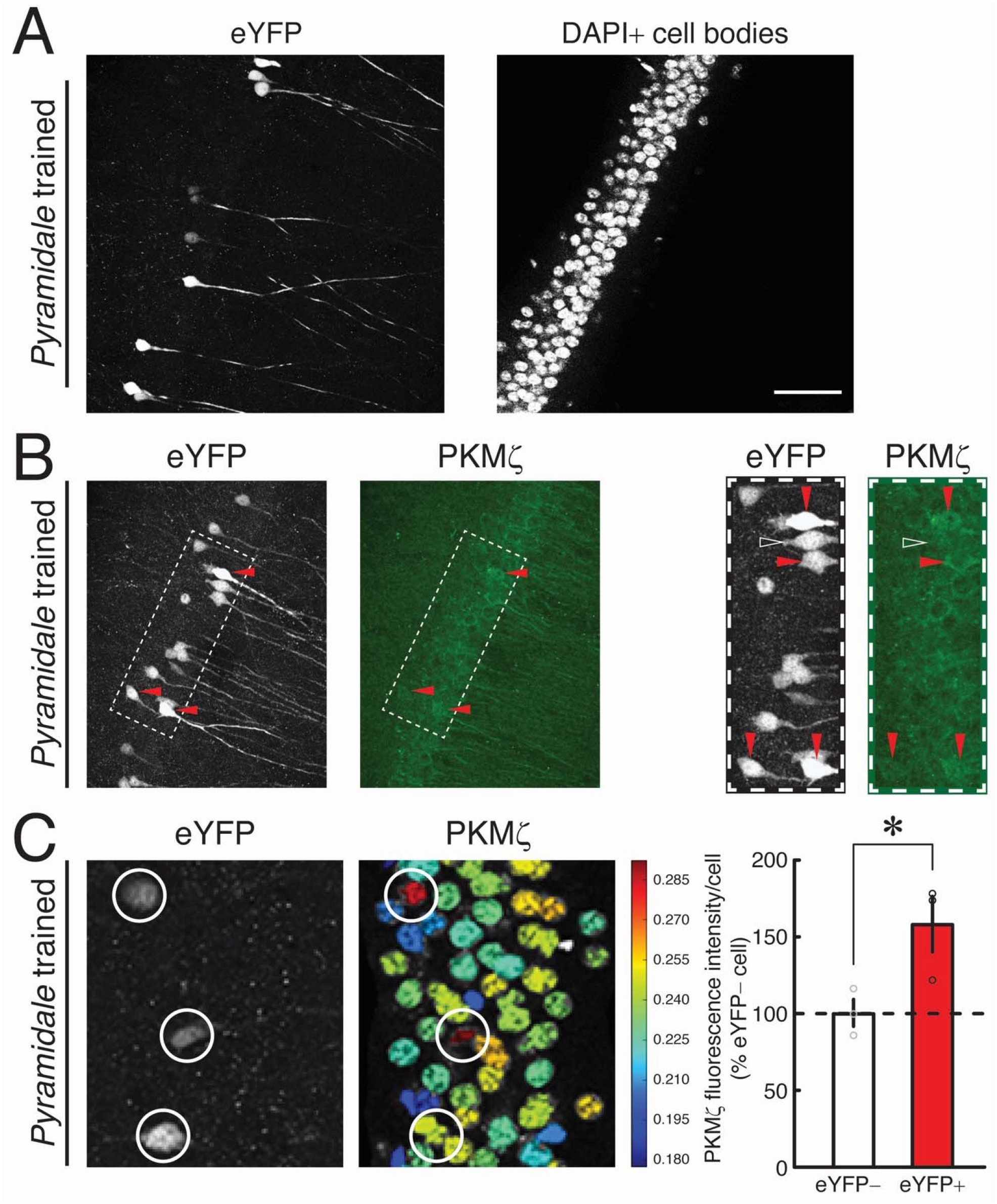
Memory-tagged CA1 pyramidal cells express more PKMζ than non-tagged cells during memory storage. To label cells active during memory retention in *Arc*Cre-eYFP mice, 1 day after training 4-OH-tamoxifen is injected 30 min before a 10-min retention test with no shock. The mice are sacrificed 7 days after the retention test, and fluorescence microscopy for eYFP+ and immunohistochemistry for PKMζ performed. (A) Left, representative image of eYFP+ neurons in CA1 pyramidal cell layer show a minority of neurons are labeled. Right, cell bodies of CA1 pyramidal cells defined by central DAPI nuclear staining. (B) eYFP+ cells in hippocampi of trained mice contain abundant PKMζ. Left panels, representative CA1 pyramidal cell layer showing eYFP and PKMζ expression. Right panels, enlarged images of the rectangles shown at left. Red arrowheads show examples of eYFP+ cells with abundant PKMζ. White open arrowhead shows a rare eYFP+ cell without excess PKMζ. (C) Quantification of PKMζ immunostaining in pyramidal cell bodies of trained mice shows abundant PKMζ in eYFP+ cells compared to eYFP– cells. Left, representative images of eYFP expression and quantification of PKMζ immunostaining in CA1 pyramidal cell layer (individual eYFP+ cells circled). Colormap is shown as insert at right. Right, mean ± SEM of the data shown in circles. Scale bar in A = A: 50 µm, B, left panels: 48 µm, B, right panels: 28 µm, C: 22 µm.

To assess the changes in number of eYFP+ cells with training, we also examined the number of eYFP+ cells in two control conditions: untrained mice, which were given the identical exposure to the rotating arena and injected at the equivalent time, but not shocked, and mice that never left the home cage, but were injected with 4-OH-tamoxifen at the equivalent times as untrained and trained mice (Supplementary Fig. 5). Because the eYFP signal can be detected as early as 1 day after 4-OH-tamoxifen injection and reaches stable levels after 5 days (data not shown), all mice remained in the isolation chamber for 7 days after 4-OH-tamoxifen injection prior to brain extraction.

### ArcCreER^T2^ x eYFP mice: IHC

Whole brain slices (250 μm) were fixed with 4% PFA and rinsed with PBS, blocked with 5% normal goat serum, permeabilized with 0.1% Triton, and incubated with primary antibodies against eYFP and PKMζ proteins (chicken anti-GFP antibody [1:1000], Abcam, and the rabbit anti-PKMζ antisera [1:5000], and fluorescence-tagged secondary antibodies, Alexa Fluor® 594 AffiniPure Goat Anti-Chicken IgY [1:500], and Alexa Fluor® 680 AffiniPure Goat Anti-Rabbit IgG [1:500], both from Jackson ImmunoResearch). Slices were then mounted with DAPI Fluoromount-G (Southern Biotech) on cover-slides (FisherBrand) and maintained at −20 °C and protected from light until confocal imaging.

Confocal microscopy imaging was performed using an Olympus FluoView FV1000 Confocal Laser Scanning Biological Microscope built on the Olympus IX81 Inverted Microscope. All imaging sessions pertaining to the same study used the same parameters (e.g.,laser power, contrast, and brightness). Confocal images (4X-60X magnification and Z-stacks of areas of interest) were exported in TIFF and JPG formats, transformed into gray-scale images, and analyzed using imaging software tools (ImageJ, version 1.50i; https://imagej.nih.gov/ij/; Rasband, W.S., ImageJ, U. S. National Institutes of Health, Bethesda, Maryland, USA, 1997-2018) and CellProfiler Analyst (CPA) (www.cellprofiler.org, (Dao *et al.*, 2016)). Image J identified and quantified eYFP-expressing cells within areas of interest. CPA quantified PKMζ expression in individual cells. CPA was first trained to recognize DAPI-containing nuclei as individual cell elements and to profile a somatic area around each one of them based on the eYFP signal from fully filled cells (false positives tested the accuracy of somatic profiling, data not shown). CPA was then trained to automatically recognize and quantify somatic PKMζ signal intensity from these cell elements in eYFP+ and eYFP– cells.

### Expansion microscopy

The density of ChR2-eYFP expression in neuropil made it difficult to evaluate immunohistochemically identified PKMζ puncta in single dendrites, and labeling with eYFP alone fills neuronal cell bodies but not the entirety of their processes (data not shown). We therefore used expansion microscopy (ExM) to isometrically enlarge the tissue sample 4.5 times (Fig. 4C). *Arc*CreER^T2^ x ChR2-eYFP mice were housed in a separate isolation room in a fresh cage (2 mice/cage) in continuous darkness 1-2 days before a pretraining trial to prevent unnecessary disturbances and reduce off-target labeling. One day after the pretraining trial, three 30-min training trials were conducted with a 2.5 h intertrial interval. The next day, mice were injected intraperitoneally with 4-OH-tamoxifen (∼200 µl of 10 µg/µl, 0.01% of body weight), and administered a fourth 30 min place avoidance training 5 h later. Following the behavioural task, mice were housed in the isolation room for the next 3 days. The mice were then taken out of isolation and returned to the normal colony room. After 2 days in the colony room, memory retention was tested (data not shown), and the mice were transcardially perfused with ice-cold saline and then 4% PFA, and the brains extracted and post-fixed in 4% PFA. Thirty µm slices through the dorsal hippocampus were cut on a vibratome and treated for IHC, as described above.

The slices were expanded (Chen *et al.*, 2015), starting with the anchoring step by using Acryloyl-X SE (6-((acryloyl)amino)hexanoic acid; 1 g/ml dissolved in dimethyl sulfoxide) stock solution overnight, followed by the gelation step, in chambers constructed by sandwiching the slice between a glass slide and coverslip with coverslip spacers.

The gelling solution was freshly prepared on ice a few minutes before use by mixing a monomer solution, consisting of: 1) sodium acrylate (8.6 g/100ml), acrylamide (2.5 g/100ml), N,N-methylenebisacrylamide (0.15 g/100ml), NaCl (11.7 g/100ml), PBS (1x), and dH2O, 2) 4-hydroxy-2,2,6,6-tetramethylpiperidin-1-oxyl stock solution (0.5 mg/100 ml), 3) tetramethylethylenediamine stock solution (10 g/100 ml), and 4) ammonium persulfate stock solution (10 g/100 ml) in a 47:1:1:1 proportion. Each tissue section was rapidly immersed in 600 µl of pre-chilled gelling solution in an Eppendorf tube and incubated for 25 min at 4 °C. The tissue sections were transferred to gelation chambers that were constructed by sandwiching the slice between a glass slide and coverslip with coverslip spacers and incubated for 2 h at 37 °C. The tissue sections were removed from the gelation chambers, and excess gel trimmed using a razor blade. The tissue was placed in a well for digestion.

The digestion buffer stock was prepared and stored at 4 °C for no longer than two weeks before use. A Triton-X (0.5 g/100 ml), EDTA (0.027 g/100 ml), Tris (5 ml/100 ml [pH 8]), NaCl (4.67 g/100 ml) and dH2O 100 ml solution was prepared. Proteinase K (BioLabs, 800 units/ml) was added to the digestion buffer stock immediately before use at 1:100 dilution. The tissue sections were allowed to digest for 20 min at room temperature without shaking, rather than overnight as previously described (Chen *et al.*, 2015). The digestion was terminated by replacing the digestion buffer with cold PBS. The tissue was stored overnight at 4 °C, followed by expansion by gradually adding ddH2O. The gradual water increase prevented tissue bending. The tissue was imaged using the inverted confocal microscope, as described above.

### Statistics

Statistica software performed all statistical analyses (StatSoft, OK). Paired Student’s *t* tests compared the average of 5 min of fEPSP responses before tetanization and 120 min after tetanization in the LTP experiments. Multi-level comparisons were performed by one-way ANOVA, and multi-factor comparisons by ANOVA with repeated measures, as appropriate. Subscripts report the degrees of freedom for the critical *t* values of the *t* tests and the *F* values of the ANOVAs. Fisher’s least significant difference (LSD) tested post-hoc multiple comparisons. After PKMζ was shown to increase in CA1 1 month after training (Fig. 3), one-tailed Student *t* tests evaluated the hypotheses that training increases PKMζ, eYFP, and their overlap (Fig. 5). Statistical significance is accepted at *P* < 0.05.

## Results

### Persistent increases in PKMζ during LTP maintenance

Persistent changes in the amount and distribution of PKMζ during late-LTP were first examined in hippocampal slices (Fig. 1A). Tetanization of Schaffer collateral/commissural fibers in CA3 induced LTP of fEPSPs recorded in CA1 *stratum radiatum* for 2 h (mean responses of the 5 min pre-tetanization compared to 5 min 2 h post-tetanization; n = 4, *t*_3_ = 8.70, *P* = 0.003). To determine the level of non-specific staining not due to PKMζ, immunostaining was compared between hippocampi from wild-type mice and PKMζ-null mice (Tsokas *et al.*, 2016) (Fig. 1B). PKMζ-null mice show minimal background immunostaining (Fig. 1B, left) that is equivalent to the low level of immunostaining in *alveus* of wild-type mice (Hernandez *et al.*, 2014) (Fig. 1B, middle), which was set as the level of background.

PKMζ-immunostaining was then compared between wild-type slices 2 h post-tetanization to control slices from the same hippocampus that received equivalent test stimulation but not tetanization (Fig. 1B, C, Supplementary Fig. 1). Low magnification images of the slices show large PKMζ increases in CA1c adjacent to the stimulating electrode that diminish towards CA1a adjacent to the subiculum, which is consistent with the projections of the Schaffer collateral fibers stimulated in CA3a (Amaral & Witter, 1989; Takacs *et al.*, 2012) (Supplementary Fig. 1). No changes are detectable in CA3 or dentate, which further supports the necessity of Schaffer collateral stimulation. We analyzed the area under the mean of PKMζ-immunointensity profiles drawn at the border between CA1b and c by ANOVA with repeated measurement (Fig. 1B, C). There is a significant main effect of treatment (untetanized and LTP maintenance, *F*_1,6_ = 25.58, *P* = 0.002), subfield (*strata oriens, pyramidale, radiatum*, and *lacunosum-moleculare, F*_3,18_ = 60.04, *P* < 0.0001), and their interaction (*F*_3,18_ = 3.51, *P* = 0.04) (Fig. 1C). Compared to untetanized slices, PKMζ intensities increase in LTP slices in *stratum pyramidale* and in dendritic *strata oriens* and *radiatum*, both of which receive projections from Schaffer collateral fibers stimulated in CA3a (Amaral & Witter, 1989; Takacs *et al.*, 2012) (post-hoc, n’s = 4, *P* = 0.002, *P* = 0.006, *P* = 0.00003, respectively). In contrast, PKMζ intensities show no change in *stratum lacunosum-moleculare*, which does not receive projections from the Schaffer collateral axons (*P* = 0.124).

### Persistent increases in PKMζ during long-term storage of spatial memory

We next examined the amount and distribution of PKMζ in the hippocampal formation during spatial long-term memory storage. The training protocol consists of a 30-min pretraining period, followed by three 30-min training trials with 2 h intertrial intervals, and a retention test 1 day later (Fig. 2A). Untrained animals were placed in the apparatus for equivalent periods of time as trained animals, but received no shock (Fig. 2A). Therefore, the only physical difference in the environment for trained and untrained mice is the 500-ms shock during each training trial (< 1% of the total training experience). Corticosterone levels do not differ between trained and untrained mice (Lesburgueres *et al.*, 2016), but the trained mice express the conditioned response whereas the untrained mice do not and thus serve as a control group. The time to first entry into the shock zone, a measurement of the conditioned response, shows significant effects of training phase (*F*_2,28_ = 12.62, *P* = 0.00012), treatment (control and trained) (*F*_1,14_ = 48.99, *P* < 0.0001), as well as their interaction (*F*_2,28_ = 13.46, *P* < 0.0001). Retention performance is significantly different by post-hoc test (*P* < 0.0001; control, n = 10, trained, n = 6).

PKMζ expression levels were examined by quantitative immunoblot one day after mice formed an active place avoidance memory (Fig. 2B). Immunoblots of mouse hippocampus 1 day after active place avoidance training show persistent increases in PKMζ compared to untrained control mice (*t*_14_ = 2.47, *P* = 0.027; untrained, n = 10, trained, n = 6). These data agree with previous work examining rat hippocampus 1 day after place avoidance training (Hsieh *et al.*, 2017).

We then examined the hippocampal formation of trained and untrained mice by immunohistochemistry (Fig. 2C, D; Supplementary Fig. 2). Inspection of images from trained mice show large increases in PKMζ in hippocampus subfield CA1 compared to untrained mice. In contrast, increases in PKMζ are less obvious in dentate gyrus (DG) and subfield CA3. The hippocampal formation shows increased PKMζ in overlying somatosensory and parietal association neocortex, at the superficial as well as the deep layers, but not retrosplenial cortex or in thalamus.

Quantitation of PKMζ-immunostaining reveals training is associated with increased PKMζ in CA1, compared to CA3 or DG (Fig. 2D). ANOVA reveals the main effects of treatment (untrained and trained, *F*_1,10_ = 8.20, *P* = 0.017), region (CA1, DG, and CA3, *F*_2,20_ = 47.00, *P* < 0.0001), and interaction between treatment and region (*F*_2,20_ = 4.97, *P* < 0.018). Post-hoc tests show that compared to the untrained control group, PKMζ in the trained group increases in CA1, but not DG or CA3 (n’s = 6, *P* < 0.0002, *P* = 0.45, *P* = 0.47, respectively). We therefore subsequently focused on characterizing the changes of PKMζ in CA1 in spatial memory.

To examine the extent to which experience in the place avoidance apparatus without shock contributes to the increase in PKMζ in CA1, we compared 3 groups: trained mice, untrained control mice placed in the apparatus without shock, and home-caged mice (Supplementary Fig. 3A). Analysis of behavior comparing the untrained and trained groups by ANOVA reveals significant effects of training phase (pretraining, the end of training, and retention, *F*_2,22_ = 19.69, *P* < 0.0001), treatment (untrained and trained, *F*_1,11_ = 29.49, *P* = 0.0002), and their interaction (*F*_2,22_ = 17.42, *P* < 0.0001). The 1-week retention performance is significantly different between groups (post-hoc test, *P* < 0.0001; untrained, n = 6, trained, n =7). PKMζ intensities in CA1 from the 3 groups were analyzed by ANOVA, which shows a treatment main effect (*F*_2,17_ = 5.16, *P* = 0.018; home-caged, n = 7, untrained, n = 6, trained, n = 7). Post-hoc tests reveal higher CA1 PKMζ intensity in the trained group than the untrained and home-caged groups (*P* = 0.033, *P* = 0.007, respectively), and no statistical difference between untrained and home-caged groups (*P* = 0.54; Supplementary Fig. 3B, C).

### Persistent increases in PKMζ in 1-month remote spatial memory storage

We next examined whether PKMζ remains increased when spatial memory has been stored for 1 month. To produce a robust hippocampus-dependent spatial memory lasting 1 month, we conditioned mice strongly by giving 3 training trials and then repeating the protocol 10 days later (Pastalkova *et al.*, 2006; Hsieh *et al.*, 2017) (Fig. 3A). Behavioral testing shows significant effects of training phase (pretraining, the end of training, and retention, *F*_2, 32_ = 7.07, *P* = 0.0029), treatment (untrained and trained, *F*_1, 16_ = 26.29, *P* < 0.0001), and their interaction (*F*_2, 32_ = 7.29, *P* = 0.0025). Retention testing 30 days after the last training trial shows a significant difference between the untrained and trained groups (post-hoc test, *P* = 0.0018; untrained control, n = 11, trained, n = 7). Immunoblotting of mouse hippocampus shows increases in PKMζ 1 month after the last training session, as compared to untrained controls (*t*_10_ = 2.65, *P* = 0.024, n’s = 6) (Fig.3B), that agrees with our previous work on rat hippocampus (Hsieh *et al.*, 2017).

Immunohistochemistry reveals that the increases in PKMζ lasting 1 month localize to CA1, similar to what was observed 1 day or 1 week after training. Because the strong conditioning made 1-month memory likely, we examined both mice that were tested for 1-month memory retention and mice that received similarly strong conditioning but were not tested.

PKMζ increases occur both in mice that were tested (*t*_6_ = 2.48, *P* = 0.048, n’s = 4) (Fig. 3C, D), and mice that were not tested (*t*_4_ = 3.73, *P* = 0.020, n’s = 3) (Supplementary Fig. 4). Inspection outside the hippocampal formation reveals PKMζ increased in deep layers of overlying somatosensory neocortex, as well as in retrosplenial cortex, but not in superficial layers as was observed a day after training (Fig. 3C).

### Persistent increases in PKMζ in subcellular compartments of active neuronal ensembles following 1-month memory storage

The persistent increases in PKMζ are hypothesized to occur preferentially in neuronal ensembles that participate in long-term memory storage. We tested this notion by examining PKMζ in mice engineered to induce permanent fluorescent tags in cells that have undergone plasticity-dependent activation of the promoter for the immediate early gene *Arc*. We examined *Arc*CreERT2 x ChR2-eYFP mice, the result of a cross between an *Arc*CreER^T2^ mouse that expresses Cre-recombinase under transcriptional control of the promoter for *Arc*, and an R26R-STOP-floxed-ChR2-eYFP mouse. The CreER^T2^ allele expresses a Cre-recombinase that translocates to the nucleus following tamoxifen treatment. Expression and translocation of the Cre-recombinase removes a floxed STOP codon, allowing synthesis of the reporter protein ChR2-eYFP. Thus, after administration of the rapidly acting 4-OH-tamoxifen to *Arc*CreERT2 x ChR2-eYFP mice, Cre recombinase is specifically expressed in cells with activated *Arc* transcription (Srinivas *et al.*, 2001; Denny *et al.*, 2014). Cells that activate the *Arc* promoter, which increase in number during training or memory retention testing, are permanently marked and can be examined during long-term memory storage (Srinivas *et al.*, 2001; Denny *et al.*, 2014). Because ChR2-eYFP is membrane-bound, the compartments of CA1 pyramidal cells that lie within different *strata* can be independently examined. Label is detected in compartments of ChR2-eYFP-positive principal cells of CA1 (for clarity referred to as “eYFP+ cells”) in *strata pyramidale, radiatum*, and *lacunosum-moleculare* (Fig. 4A, B). In parallel experiments using expansion microscopy of tissue from CA1 radiatum, the eYFP outlining of membranes can be observed extending into dendritic spines where PKMζ may be present (Hernandez *et al.*, 2014) (Fig. 4C).

We then set out to evaluate the central hypothesis of this work that the distribution of PKMζ within compartments of active neuronal ensembles reveals specific sites of the molecular mechanism of LTP maintenance in hippocampus during spatial long-term memory storage. We tested whether PKMζ persistently increases in compartments of memory training-tagged eYFP+ cells, as compared to eYFP+ cells in control mice that had similar exposure to the apparatus without training. We examined a 1-month active place avoidance memory that allowed sufficient time for the membrane-bound eYFP to distribute throughout the cell, and in which differences between the trained and untrained control mice are expected to be related to memory and not a consequence of recent conditioning or active consolidation. The training protocol that repeats the conditioning after a 10-day interval raises the possibility that interpreting the memory-related tagging would be complicated by memory reconsolidation. Therefore, cells were tagged by eYFP soon after mice had learned the conditioned response in a long-term memory protocol that induces memory retention for 1 month with a minimum of conditioning (Fig. 5A). Trained and untrained *Arc*CreERT2 x ChR2-eYFP mice receive four 30-min trials on day 1 with a 2 h intertrial interval. The first trial is a pretraining trial with no shock, followed by 3 training trials with shock. The mice are returned to the isolation chamber after each trial. Thirty minutes before the third training trial, when the task is familiar and the trained mice demonstrate strong conditioned active place avoidance memory, all mice are injected with 4-OH-tamoxifen and returned to their home cage in the isolation chamber until the third training trial. One month after the training, the mice are returned to the arena for 10-min without shock to evaluate memory retention. The mice are sacrificed, and their brains prepared for IHC.

An analysis of behavior reveals that memory is formed during training and persists for 1 month. Two-way ANOVA shows main effects of treatment (untrained and trained, n’s = 4, *F*_1,6_ = 113.5, *P* < 0.0001), training phase (pretraining, the end of training, and retention, *F*_2,12_ = 7.83, *P*< 0.01), and their interaction (*F*_2,12_ = 7.32, *P* < 0.01). Post-hoc tests show that during the pretraining trial, trained and untrained mice do not differ on time to enter the shock zone after being placed on the arena (*P* = 0.99). By the third and last training trial, the trained mice remember to delay entering the shock zone for several minutes before any shocks were given on the trial (*P* = 0.000019). This conditioned response remains 1 month later on the retention test, during which no shocks are given (*P* = 0.02; Fig. 5A).

We evaluated whether PKMζ selectively increases in the memory-tagged eYFP+ cells, and, furthermore, whether PKMζ preferentially increases in eYFP+ cells at *stratum radiatum* compared to *stratum lacunosum-moleculare*, as predicted by evidence that CA3→CA1 synapses preferentially process spatial memory (Brun *et al.*, 2002; Lisman, 2005; Colgin *et al.*, 2009; Pavlowsky *et al.*, 2017; Choi *et al.*, 2018; Dvorak *et al.*, 2018). For eYFP and PKMζ IHC, ROIs centered on *strata pyramidale, radiatum*, and *lacunosum-moleculare* were examined (Fig. 5B). The raw integrated density (defined as the sum of the values for all pixels) of the ROI expressing the fluorescent label was measured for the green (eYFP), red (PKMζ), and yellow (overlap) volume of pixels. These measures account for the locations of the signal as well as their intensity.

As our primary goal is to evaluate whether training increases PKMζ in the memory-tagged eYFP-expressing neurons, it is important to first distinguish between an increase in overlap between eYFP and PKMζ following training that might possibly result because of general increases in eYFP and PKMζ (the null hypothesis, HO), or, alternatively, because PKMζ selectively increases in the eYFP+ cells that are an enriched sample of memory-associated neurons (H1). HO predicts the independence of expression of eYFP and PKMζ, and thus that the probability of PKMζ given eYFP [p(PKMζ | EYFP)] equals the product of p(PKMζ) x p(eYFP). In contrast, H1 predicts the conditional probability of PKMζ given eYFP will be greater than the product of their individual probabilities.

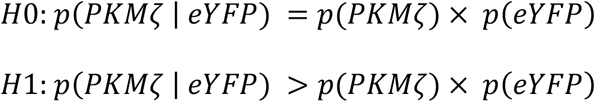

In both trained and untrained mice the conditional probability is greater in almost all mice in *strata pyramidale, radiatum*, and *lacunosum-moleculare* compartments (*t*_23_ = 5.08, *P* < 0.00001) (Fig. 5C), supporting the hypothesis that experience increases PKMζ selectively in eYFP+ cells. One-tailed Student *t* tests evaluated the hypotheses that training increases the hippocampal expression of eYFP, PKMζ, and their overlap (Fig. 5D). Labeling intensities for eYFP, PKMζ, and their overlap (normalized by the combined eYFP and PKMζ signal) significantly increased in the trained group in *stratum pyramidale* for eYFP (*t*_7_ = 2.007, *P* = 0.04), for PKMζ (*t*_7_ = 2.12, *P* = 0.03), and for their overlap (*t*_7_ = 2.54, *P* = 0.02). A similar finding was seen in *stratum radiatum* for eYFP (*t*_7_ = 2.67, *P* = 0.01), for PKMζ (*t*_7_ = 2.85, *P* = 0.01), and for their overlap (*t*_7_ = 3.34, *P* = 0.007). In contrast, no group differences were found in *stratum lacunosum-moleculare* (eYFP: *t*_7_ =1.47, *P* = 0.09; PKMζ: *t*_7_ = 1.23, *P* = 0.13, and overlap: *t*_7_ = 0.02, *P* = 0.49).

We examined the overlap between PKMζ and eYFP in more detail. Separate populations of memory-allocated cells activate distinct plasticity-related immediate early genes in addition to *Arc*. These genes include *Fos, Fosl2, Egr1, Jun, Homer1, Npas4*, and others (Denny *et al.*, 2014; Harris *et al.*, 2020; Sun *et al.*, 2020). Therefore, only a fraction of memory-activated cells are likely to be labeled by eYPF. Furthermore, we presume that in addition to the expression of PKMζ in eYFP+ cells, PKMζ is also expressed independently to maintain other memories. Consequently, we computed the Manders coefficients M1 and M2 to rigorously compare the overlap between eYFP and PKMζ signals in the hippocampi of trained and untrained mice (Manders *et al.*, 1993). M2 (eYFP&PKMζ / eYFP) computes the proportion of the eYFP signal that colocalizes with the PKMζ signal. Accordingly, if memory training recruits cells, some of which express eYFP, and all recruited cells express PKMζ, then M2 will be greater in tissue from trained than untrained mice. In contrast, M1 (PKMζ&eYFP / PKMζ) estimates the proportion of the PKMζ signal that colocalizes with the eYFP signal. How memory training should change M1 is not obvious if PKMζ is expressed by memories formed prior to the place avoidance conditioning, and also after training in memory-activated cells that express immediate early genes other than *Arc*.

M2 (eYFP&PKMζ / eYFP) is significantly increased in the trained group in *stratum pyramidale* (*t*_7_ = 2.16, *P* = 0.03) and *stratum radiatum* (*t*_7_ = 2.89, *P* = 0.01), as predicted by memory-induced persistent PKMζ expression, whereas no difference is observed in *stratum lacunosum-moleculare* (*t*_7_ = 0.051, *P* = 0.44) (Fig. 5D). Thus, the M2 analysis identifies that memory training increases PKMζ expression in the *stratum radiatum* of eYFP+ cells. In contrast, whereas both the memory-training and control experiences induce expression of eYFP in neurons, there was no group difference in M1 (PKMζ&eYFP / PKMζ) in *stratum pyramidale* (*t*_7_ = 0.52, *P* = 0.62), *stratum radiatum* (*t*_7_ = 1.38, *P* = 0.21), or *stratum lacunosum-moleculare* (*t*_7_ = 2.31, *P* = 0.06.) Thus, whereas both the memory and control experiences induce eYFP expression, the proportion of the PKMζ-expressing cells that also express eYFP does not change between the memory-training and control experiences. This result indicates that PKMζ is expressed in cells in addition to those with *Arc*-promoter activation of eYFP expression, and this proportion does not differ between the two experiences.

Collectively, these findings strongly support the hypothesis that neurons that are active during the expression of memory also persistently increase PKMζ levels for at least 1 month. Furthermore, although activation of *Arc* is sufficient to label memory-related cells that show persistent increases in PKMζ, the M1 findings suggest that the *Arc* promoter-labeling represents only a subset of memory-activated cells (Denny *et al.*, 2014; Harris *et al.*, 2020; Sun *et al.*, 2020).

We also measured the percent area of the ROI that expresses the fluorescent label. Again, the area labeled with eYFP, PKMζ, and their overlap increases in the trained group in *stratum pyramidale* (eYFP: t7 = 2.18, *P* = 0.03; PKMζ: *t*_7_ = 2.44, *P* = 0.02; and overlap: *t*_7_ = 2.19, *P* = 0.03), and in *stratum radiatum* (eYFP: *t*_7_ = 2.59, *P* = 0.02; PKMζ: *t*_7_ = 1.983, *P* = 0.04; and overlap: *t*_7_ = 2.274, *P* = 0.03). No significant group differences are observed in *stratum lacunosum-moleculare* (eYFP: *t*_7_ = 1.77, *P* = 0.43; PKMζ: *t*_7_ = 1.68, *P* = 0.11; overlap: *t*_7_ = 0.31, *P* = 0.38).

In addition to comparing PKMζ expression in eYFP+ cells of trained mice and untrained mice, we also examined differences in PKMζ expression between eYFP+ and eYFP– cells within the same hippocampi of trained mice. We took advantage of our findings that showed training increases PKMζ in *stratum pyramidale* (Figs. 2, 3, 5, Supplementary Figs. 2, 3, 4) and estimated the centers of the cell bodies of eYFP+ and eYFP– cells using nuclear DAPI-staining (Fig. 6A). To ensure that the cell bodies of the memory-tagged neurons are completely filled with fluorescent marker, we used *Arc*CreER^T2^ x eYFP mice, in which the plasticity-dependent marker is the cytosolic protein, eYFP.

PKMζ expression was hypothesized to increase in eYFP+ cells compared to eYFP– cells during long-term memory retention. This hypothesis was tested by training mice and then injecting 4-OH-tamoxifen 1 day later, 30 min prior to a retention test without shock. The animals are sacrificed 1 week after the 4-OH-tamoxifen injections, allowing time for stable somatic eYFP expression (Srinivas *et al.*, 2001; Denny *et al.*, 2014). The percent of CA1 eYFP+ cells in the trained mice is 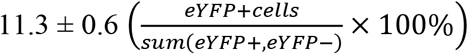, in line with prior results (Denny *et al.*, 2014). To determine whether the percent of eYFP+ cells in CA1 is due to an increase induced by the retention of memory, we compared the percent of the eYFP+ cells in the CA1 of the trained mice to that of untrained (4.2 ± 0.3%) and home-caged mice (2.8 ± 0.2%) (Supplementary Fig.5). One-way ANOVA shows the main effect of treatment (trained, untrained, and home-caged, *F*_2,12_ = 125.2, *P* < 0.00001), and post-hoc tests show that the percent of eYFP+ cells of the trained group is higher than both the untrained and home-cage groups (n’s = 5, *P*’s < 0.00001), and the percent in the untrained group is also higher than in the home-caged group (n’s = 5, *P* = 0.043). The amount of PKMζ in the soma of eYFP+ cells and eYFP– cells was then compared within the same CA1 after training (Fig. 6B, C). The results reveal PKMζ is more abundant in eYFP+ cells than in eYFP– cells (n’s = 3, *t*_4_ = 2.85, *P* = 0.046). Thus, PKMζ expression in eYFP+ cells is both more than in neighboring eYFP– cells in trained mice (Fig. 6) and more than in the eYFP+ cells of untrained mice (Fig. 5).

## Discussion

This study shows that the amount and distribution of hippocampal PKMζ persistently increases after LTP induction and spatial conditioning. These persistent increases provide strong evidence that PKMζ both maintains synaptic potentiation and the storage of long-term memory. During LTP maintenance, increased PKMζ persists in CA1 for at least 2 h in hippocampal slices in projection fields that had received strong afferent stimulation (Fig. 1, Supplementary Fig. 1).

During spatial long-term memory, increased PKMζ persists for at least 1 month in the somatic and *stratum radiatum* dendritic compartments of CA1 neuronal ensembles that activated plasticity-dependent gene expression during the initial stage of memory storage (Figs. 3, 4, 5). To our knowledge, these are the most long-lasting changes in expression of a protein in hippocampus-dependent memory. Increases in PKMζ are sufficient to potentiate synaptic transmission, and suppression of PKMζ activity reverses the maintenance of late-LTP and hippocampus-dependent memory up to 1 month after training (Ling *et al.*, 2002; Pastalkova *et al.*, 2006; Tsokas *et al.*, 2016; Wang *et al.*, 2016). Taken with the results of this study, we conclude that the loci of persistently increased PKMζ represent sites of the endogenous molecular mechanism of LTP maintenance that sustain spatial long-term memory.

In LTP, PKMζ persistently increases in the projection fields of the tetanically stimulated afferent fibers (Figs. 1B, C, Supplementary Fig. 1). LTP was induced in hippocampal slices using stimulating electrodes placed in *stratum radiatum* of CA3a, the CA3 subregion closest to CA1. The CA3a Schaffer collateral fibers synapse on CA1c pyramidal cells on apical dendrites in *stratum radiatum* and basal dendrites in *stratum oriens* (Amaral & Witter, 1989; Takacs *et al.*, 2012). These experiments were performed in transverse slices from dorsal hippocampus where CA3a Schaffer collateral projections to *stratum oriens* are particularly numerous (Amaral & Witter, 1989). Both CA1c *strata radiatum* and *oriens* show persistent increases in PKMζ during LTP maintenance, whereas regions with fewer or no CA3a projections, such as CA1c *stratum lacunosum-moleculare* or CA1a, do not (Fig. 1B, C, Supplementary Fig. 1).

Mechanisms sustaining increased PKMζ in LTP might include local synthesis of PKMζ from its dendritic mRNA, similar to the local synthesis underlying increased dendritic CaMKII seen 30 min after tetanization in LTP (Ouyang *et al.*, 1997). Alternatively, newly synthesized PKMζ may translocate to recently activated postsynaptic sites (Sajikumar *et al.*, 2005; Palida *et al.*, 2015). PKMζ also persistently increases in CA1 pyramidal cell bodies (Figs. 4, 6). Somatic increases might represent sites of non-synaptic actions of PKMζ important for long-term neuronal plasticity and memory maintenance, particularly the epigenetic control of gene transcription in nuclei (Hernandez *et al.*, 2014; Ko *et al.*, 2016). High basal levels of PKMζ are also seen in *stratum lacunosum-moleculare* (Fig. 1B), which might also be sustained by ongoing local synthesis or post-translational compartmentalization. Such basal levels of PKMζ may represent PKMζ formed by long-term memory storage of prior experiences.

PKMζ persistently increases in CA1 from 1 day to at least 1 month after spatial conditioning. Memory retention testing is not needed for these very long-term increases, suggesting that increased PKMζ activity underlies both maintenance and expression of long-term memory (Fig. 3, Supplementary Fig. 4) (Sacktor & Fenton, 2018). In contrast to LTP in which PKMζ preferentially increases in the projections fields of the tetanically stimulated fibers (Supplementary Fig. 1), spatial conditioning increases PKMζ throughout CA1 (Figs. 2, 3, Supplementary Figs. 3, 4). This localization of persistent increases in PKMζ to the output neurons of the hippocampal formation in this spatial task is analogous to similar increases in the output neurons of motor cortex during long-term retention of a reaching task (Gao *et al.*, 2018). Neocortical PKMζ also appears to increase in active place avoidance memory at both superficial and deep layers at the earlier time point, but less so at the superficial layers in remote memory, while increasing at deep layers of retrosplenial cortex during remote memory, but not at the earlier time point (Wesierska *et al.*, 2009). The changes in PKMζ outside the hippocampus will require future study, and highlight that there are system-level dynamics underlying the location of persistent PKMζ expression and perhaps LTP maintenance across time scales of days and weeks. Using the present approach will also be important for determining to what extent the observed changes are conserved for other memories that rely on PKMζ for their persistence (Serrano *et al.*, 2008; Tsokas *et al.*, 2016).

During spatial long-term memory retention, both the expression of PKMζ and the number of *Arc* promoter-driven, memory-tagged cells increase more in CA1 than in CA3 or DG. (Fig. 2C, D, Supplementary Fig. 5). The paucity of PKMζ increases and *Arc*-promoter activation in CA3 contrasts with the prominent role of recurrent collaterals of CA3 pyramidal cells in theories of hippocampal memory storage (McNaughton & Morris, 1987; Jensen & Lisman, 1996; Rolls, 1996; Papp *et al.*, 2007). Kinases or signaling molecules other than PKMζ, such as CaMKII (Ouyang *et al.*, 1997; Lisman, 2017) or other PKCs (Hsieh *et al.*, 2017), may play roles in long-term memory storage in CA3. Alternatively, because spatial conditioning induces fewer traces of PKMζ or memory-tagged cells in CA3, this region may play a role in storing short-term rather than long-term memories for active place avoidance (Gilbert & Kesner, 2006). Long-term information might also be stored in only a few eYFP+ cells with PKMζ in CA3 or DG. This possibility may be the case in DG, where inspection suggests the possibility of increased PKMζ and *Arc* promoter activation in its suprapyramidal blade (Fig. 2D, Supplementary Fig. 5), and where there may also be preferential roles of the immediate early genes *Fos* and *Npas4* for labeling the granule cells that undergo physiological changes (Sun *et al.*, 2020).

Spatial training induces more PKMζ and eYFP+ cells in CA1 compared to untrained and home-caged mice (Figs. 2, 3, 5, Supplementary Figs. 2-5). In contrast, untrained mice having similar exposure to the apparatus without shock had no more PKMζ and only small increases in eYFP+ cells compared to home-caged animals (Supplementary Figs. 3, 5). The small increase in eYFP+ cells in untrained mice may reflect that activation of immediate early genes such as *Arc* mark neural activity that induces synaptic plasticity, rather than neural activity *per se* (Vazdarjanova *et al.*, 2006). ARC expression is upregulated in CA1 during exploration of a novel environment, but attenuates after repeated exposures to the same environment (Guzowski *et al.*, 2006), which is analogous to the experience of untrained control mice in our study. In part, the relative paucity of PKMζ increases in untrained mice compared to trained mice might reflect that fewer *Arc*-promoter activated, eYFP+ cells were induced in this condition.

We used this knowledge of *Arc* activation to label memory-related pyramidal cells with eYFP at the end of a series of training trials (Fig. 5) or 1 day after training (Fig. 6), when the environment and active place avoidance or control tasks were familiar to the mice. Labeling cells with eYFP after learning had occurred allowed us to separate the process of tagging from the experience of learning the relationship between the conditioned stimuli (places) and the unconditioned stimulus (foot shock). Thus, the timing of 4-OH-tamoxifen injections in both trained and untrained control mice produced an enriched sample of tagged neurons associated with familiar recollections, rather than novel experiences. For trained mice, these recollections include the location of shock.

Remarkably, not only is PKMζ expression increased for the duration of a 1-month-old place avoidance memory, but the kinase is preferentially increased in eYFP+ cells in trained mice compared to untrained mice (Fig. 5). These results indicate that PKMζ is associated with the place avoidance memory and not merely with *Arc*-promoter activation at the time of 4-OH-tamoxifen administration. These findings are similar to effects seen with perceptual learning in primary auditory cortex in which ARC expression is associated with neuroplasticity rather than neural activity (Carpenter-Hyland *et al.*, 2010). Thus, even though both trained and untrained experiences label cells with eYFP, only training that forms long-term memory might produce long-term increases of PKMζ in the labeled cells. This hypothesis is supported by previous results showing strong place avoidance conditioning causes persistent increases of PKMζ in the hippocampus, whereas weaker conditioning produces only transient increases (Hsieh *et al.*,2017).

The PKMζ increase in CA1 pyramidal cells in 1-month-old memory occurs selectively in dendrites within *stratum radiatum*, indicating long-term storage of information received by Schaffer collateral/commissural pathways from CA3 to CA1 pyramidal cells. These increases were not seen in *stratum lacunosum-moleculare* that receives projections of the perforant pathway from EC3 to CA1 pyramidal cells. These results validate our central hypothesis that persistent increases in PKMζ reveal specific sites of the physiological maintenance mechanism of LTP within the dorsal hippocampus during spatial long-term memory storage, as predicted by prior physiological work (Brun *et al.*, 2002; Lisman, 2005; Colgin *et al.*, 2009; Pavlowsky *et al.*, 2017; Choi *et al.*, 2018; Dvorak *et al.*, 2018). In particular, the results agree with observations of synaptic potentiation of Schaffer collateral/commissural pathways, but not of temporoammonic pathways, for at least 1 month after long-term place avoidance conditioning, but only if the mouse expresses the 1-month memory (Pavlowsky *et al.*, 2017). PKMζ is present in some spines and not others within the eYFP+ dendritic compartments in *stratum radiatum* of trained animals (Fig. 4C). Because only potentiated synaptic pathways are depotentiated by PKMζ inhibition (Ling *et al.*, 2002; Sajikumar *et al.*, 2005; Pastalkova *et al.*, 2006), we speculate that PKMζ may be present selectively in potentiated spines that receive simultaneous pre- and postsynaptic activation during learning. Future work with techniques such as dual enhanced green fluorescent protein reconstitution across synaptic partners (dual-eGRASP) will be required to test this hypothesis (Choi *et al.*, 2018). In addition to spines, we observe PKMζ in dendritic shafts (Fig. 4C), in agreement with immunoelectronmicroscopy showing PKMζ both within spines and outside of spines in dendritic granules and endoplasmic reticulum, as well as more diffusely (Hernandez *et al.*, 2014). These extraspinous locations suggest additional roles for PKMζ, such as in persistent upregulation of protein synthesis in dendrites that might form positive feedback loops that maintain the sustained increases of the kinase (Westmark *et al.*, 2010; Helfer & Shultz, 2018). These observations of compartment-specific, persistent increases in PKMζ expression indicate that technologies that can selectively manipulate PKMζ function in specific somato-dendritic compartments will be needed to conduct causal experiments that elucidate their roles in maintaining long-term memory.

The sustained increases in PKMζ expression that can last for weeks might be due to positive feedback loops that persistently increase its synthesis (Westmark *et al.*, 2010; Jalil *et al.*, 2015; Helfer & Shultz, 2018) or decrease its turnover (Vogt-Eisele *et al.*, 2013). This steady-state increase is far longer than the putative half-life of PKMζ, which is likely hours to days, as suggested by PKMζ protein downregulation by shRNA (Dong *et al.*, 2015; Wang *et al.*, 2016) and inducible genetic knockdown (Volk *et al.*, 2013). Although others have argued that a relatively short protein half-life makes PKMζ an unlikely storage mechanism for very long-term memory (Palida *et al.*, 2015), our results show that this is not necessarily the case. Indeed, in primary cultures of neurons in which the PKMζ half-life has been measured, the persistent steady-state increase of PKMζ induced by chemical-LTP far outlasts the half-life of individual PKMζ protein molecules (Palida *et al.*, 2015). Our results *in vivo* suggest that these steady-state increases can reveal the physical substrate of long-term memory storage in the brain.

### Supporting Information

Additional supporting information can be found in the online version of this article.

### Data Accessibility

Research data will be made available on request to the corresponding authors.

## Acknowledgements

We gratefully acknowledge support from grants provided by NIH grants R37 MH057068 (T.C.S.), R01 MH115304 (T.C.S., A.A.F.), R01 NS108190 (P.B., T.C.S.), R01 NS105472 (A.A.F.), R21 NS091830 (J.M.A., A.A.F.), and the Garry & Sarah S. Sklar Fund (P.T.). P.T. is an Alexander S. Onassis Public Benefit Foundation Scholar. We dedicate this paper to the memories of Dr. Ned Charlton Sacktor and Andrew Carter.

## Author contributions

C.H. and P.T. contributed equally to this study, T.C.S, A.A.F., J.M.A., A.G.-P., J.B., K.K., J.C., A.C., C. J., E.L., N.S.B., C.A.D., A.I.H., P.J.B., J.E.C., C.H., and P.T. conceived the study and designed the methodology. C.H., P.T., A.G.-P., J.B., K.K., J.C., A.C., C. J., E.L. performed the experiments. C.H., P.T., A.G.-P., J.B., K.K., J.C., A.C., C.J., E.L., R.E.F.-O., L.M.R.V., A.A.F. and T.C.S. performed the analyses of the experimental data. T.C.S., P.J.B., J.M.A., and A.A.F. wrote the article with contributions and input from N.S.B., C.A.D., C.H., P.T., and A.G.-P.

## Abbreviations

CA1: *Cornu Ammonis 1* subfield
CA3: *Cornu Ammonis 3* subfield
DG: dentate gyrus
EC3: entorhinal cortex layer 3
eYFP: enhanced yellow fluorescent protein
fEPSPs: field excitatory postsynaptic potentials
IHC: immunohistochemistry
LTP: long-term potentiation
PKC: protein kinase C
PKMζ: protein kinase Mζ
SEM: standard error of mean

## References

Amaral, D.G. & Witter, M.P. (1989) The three‐dimensional organization of the hippocampal formation: a review of anatomical data. Neuroscience, 31, 571‐591.

Anderson, W.W. & Collingridge, G.L. (2007) Capabilities of the WinLTP data acquisition program extending beyond basic LTP experimental functions. J Neurosci Methods, 162, 346‐356.

Brun, V.H., Otnass, M.K., Molden, S., Steffenach, H.A., Witter, M.P., Moser, M.B. & Moser, E.I. (2002) Place cells and place recognition maintained by direct entorhinal‐hippocampal circuitry. Science, 296, 2243‐2246.

Carpenter‐Hyland, E.P., Plummer, T.K., Vazdarjanova, A. & Blake, D.T. (2010) Arc expression and neuroplasticity in primary auditory cortex during initial learning are inversely related to neural activity. Proc Natl Acad Sci U S A, 107, 14828‐14832.

Chen, F., Tillberg, P.W. & Boyden, E.S. (2015) Optical imaging. Expansion microscopy. Science, 347, 543‐548.

Choi, J.H., Sim, S.E., Kim, J.I., Choi, D.I., Oh, J., Ye, S., Lee, J., Kim, T., Ko, H.G., Lim, C.S. & Kaang, B.K. (2018) Interregional synaptic maps among engram cells underlie memory formation. Science, 360, 430‐435.

Colgin, L.L., Denninger, T., Fyhn, M., Hafting, T., Bonnevie, T., Jensen, O., Moser, M.B. & Moser, E.I. (2009) Frequency of gamma oscillations routes flow of information in the hippocampus. Nature, 462, 353‐357.

Dao, D., Fraser, A.N., Hung, J., Ljosa, V., Singh, S. & Carpenter, A.E. (2016) CellProfiler Analyst: interactive data exploration, analysis and classification of large biological image sets. Bioinformatics, 32, 3210‐3212.

Denny, C.A., Kheirbek, M.A., Alba, E.L., Tanaka, K.F., Brachman, R.A., Laughman, K.B., Tomm, N.K., Turi, G.F., Losonczy, A. & Hen, R. (2014) Hippocampal memory traces are differentially modulated by experience, time, and adult neurogenesis. Neuron, 83, 189‐201.

Dong, Z., Han, H., Li, H., Bai, Y., Wang, W., Tu, M., Peng, Y., Zhou, L., He, W., Wu, X., Tan, T., Liu, M., Wu, X., Zhou, W., Jin, W., Zhang, S., Sacktor, T.C., Li, T., Song, W. & Wang, Y.T. (2015) Long‐term potentiation decay and memory loss are mediated by AMPAR endocytosis. The Journal of clinical investigation, 125, 234‐247.

Dvorak, D., Radwan, B., Sparks, F.T., Talbot, Z.N. & Fenton, A.A. (2018) Control of recollection by slow gamma dominating mid‐frequency gamma in hippocampus CA1. PLoS Biol, 16, e2003354.

Gao, P.P., Goodman, J.H., Sacktor, T.C. & Francis, J.T. (2018) Persistent increases of PKMζ in sensorimotor cortex maintain procedural long‐term memory storage. iScience, 5, 90‐98.

Gilbert, P.E. & Kesner, R.P. (2006) The role of the dorsal CA3 hippocampal subregion in spatial working memory and pattern separation. Behavioural brain research, 169, 142‐149.

Guzowski, J.F., Miyashita, T., Chawla, M.K., Sanderson, J., Maes, L.I., Houston, F.P., Lipa, P., McNaughton, B.L., Worley, P.F. & Barnes, C.A. (2006) Recent behavioral history modifies coupling between cell activity and Arc gene transcription in hippocampal CA1 neurons. Proc Natl Acad Sci U S A, 103, 1077‐1082.

Harris, R.M., Kao, H.-Y., Alarcon, J.M., Fenton, A.A. & Hofmann, H.A. (2020) Transcriptome analysis of hippocampal subfields identifies gene expression profiles associated with long‐term active place avoidance memory. bioRxiv 2020.02.05.935759.

Helfer, P. & Shultz, T.R. (2018) Coupled feedback loops maintain synaptic long‐term potentiation: A computational model of PKMzeta synthesis and AMPA receptor trafficking. PLoS Comput Biol, 14, e1006147.

Hernandez, A.I., Blace, N., Crary, J.F., Serrano, P.A., Leitges, M., Libien, J.M., Weinstein, G., Tcherapanov, A. & Sacktor, T.C. (2003) Protein kinase Mζ synthesis from a brain mRNA encoding an independent protein kinase Cζ catalytic domain. Implications for the molecular mechanism of memory. J Biol Chem, 278, 40305‐40316.

Hernandez, A.I., Oxberry, W.C., Crary, J.F., Mirra, S.S. & Sacktor, T.C. (2014) Cellular and subcellular localization of PKMzeta. Philosophical transactions of the Royal Society of London. Series B, Biological sciences, 369, 20130140.

Hsieh, C., Tsokas, P., Serrano, P., Hernandez, A.I., Tian, D., Cottrell, J.E., Shouval, H.Z., Fenton, A.A. & Sacktor, T.C. (2017) Persistent increased PKMzeta in long‐term and remote spatial memory. Neurobiology of learning and memory, 138, 135‐144.

Jalil, S.J., Sacktor, T.C. & Shouval, H.Z. (2015) Atypical PKCs in memory maintenance: the roles of feedback and redundancy. Learning & memory, 22, 344‐353.

Jensen, O. & Lisman, J.E. (1996) Hippocampal CA3 region predicts memory sequences: accounting for the phase precession of place cells. Learning & memory, 3, 279‐287.

Ko, H.G., Kim, J.I., Sim, S.E., Kim, T., Yoo, J., Choi, S.L., Baek, S.H., Yu, W.J., Yoon, J.B., Sacktor, T.C. & Kaang, B.K. (2016) The role of nuclear PKMzeta in memory maintenance. Neurobiology of learning and memory, 135, 50‐56.

Lacagnina, A.F., Brockway, E.T., Crovetti, C.R., Shue, F., McCarty, M.J., Sattler, K.P., Lim, S.C., Santos, S.L., Denny, C.A. & Drew, M.R. (2019) Distinct hippocampal engrams control extinction and relapse of fear memory. Nat Neurosci, 22, 753‐761.

Lesburgueres, E., Sparks, F.T., O’Reilly, K.C. & Fenton, A.A. (2016) Active place avoidance is no more stressful than unreinforced exploration of a familiar environment. Hippocampus, 26, 1481‐1485.

Ling, D.S., Benardo, L.S., Serrano, P.A., Blace, N., Kelly, M.T., Crary, J.F. & Sacktor, T.C. (2002) Protein kinase Mζ is necessary and sufficient for LTP maintenance. Nat Neurosci, 5, 295‐296.

Lisman, J. (2017) Criteria for identifying the molecular basis of the engram (CaMKII, PKMzeta). Molecular brain, 10, 55.

Lisman, J.E. (2005) Hippocampus, II: memory connections. Am J Psychiatry, 162, 239.

Manders, E.M.M., Verbeek, F.J. & Aten, J.A. (1993) Measurement of co‐localization of objects in dual‐colour confocal images. J Microscopy, 169, 375‐382.

McNaughton, B.L. & Morris, R.G.M. (1987) Hippocampal synaptic enhancement and information storage within a distributed memory system. Trends Neurosci, 10, 409‐415.

Muslimov, I.A., Nimmrich, V., Hernandez, A.I., Tcherepanov, A., Sacktor, T.C. & Tiedge, H. (2004) Dendritic transport and localization of protein kinase Mζ mRNA: Implications for molecular memory consolidation. J Biol Chem, 279, 52613‐52622.

Osten, P., Valsamis, L., Harris, A. & Sacktor, T.C. (1996) Protein synthesis‐dependent formation of protein kinase Mζ in long‐term potentiation. J Neurosci, 16, 2444‐2451.

Ouyang, Y., Kantor, D., Harris, K.M., Schuman, E.M. & Kennedy, M.B. (1997) Visualization of the distribution of autophosphorylated calcium/calmodulin‐dependent protein kinase II after tetanic stimulation in the CA1 area of the hippocampus. J Neurosci, 17, 5416‐5427.

Palida, S.F., Butko, M.T., Ngo, J.T., Mackey, M.R., Gross, L.A., Ellisman, M.H. & Tsien, R.Y. (2015) PKMzeta, But Not PKClambda, Is Rapidly Synthesized and Degraded at the Neuronal Synapse. J Neurosci, 35, 7736‐7749.

Papp, G., Witter, M.P. & Treves, A. (2007) The CA3 network as a memory store for spatial representations. Learning & memory, 14, 732‐744.

Pastalkova, E., Serrano, P., Pinkhasova, D., Wallace, E., Fenton, A.A. & Sacktor, T.C. (2006) Storage of spatial information by the maintenance mechanism of LTP. Science, 313, 1141‐1144.

Pavlowsky, A., Wallace, E., Fenton, A.A. & Alarcon, J.M. (2017) Persistent modifications of hippocampal synaptic function during remote spatial memory. Neurobiology of learning and memory, 138, 182‐197.

Perusini, J.N., Cajigas, S.A., Cohensedgh, O., Lim, S.C., Pavlova, I.P., Donaldson, Z.R. & Denny, C.A. (2017) Optogenetic stimulation of dentate gyrus engrams restores memory in Alzheimer’s disease mice. Hippocampus, 27, 1110‐1122.

Rolls, E.T. (1996) A theory of hippocampal function in memory. Hippocampus, 6, 601‐620.

Sacktor, T.C. & Fenton, A.A. (2018) What does LTP tell us about the roles of CaMKII and PKMzeta in memory? Molecular brain, 11, 77.

Sacktor, T.C., Osten, P., Valsamis, H., Jiang, X., Naik, M.U. & Sublette, E. (1993) Persistent activation of the ζ isoform of protein kinase C in the maintenance of long‐term potentiation. Proc Natl Acad Sci USA, 90, 8342‐8346.

Sajikumar, S., Navakkode, S., Sacktor, T.C. & Frey, J.U. (2005) Synaptic tagging and cross‐tagging: the role of protein kinase Mζ in maintaining long‐term potentiation but not long‐term depression. J Neurosci, 25, 5750‐5756.

Serrano, P., Friedman, E.L., Kenney, J., Taubenfeld, S.M., Zimmerman, J.M., Hanna, J., Alberini, C., Kelley, A.E., Maren, S., Rudy, J.W., Yin, J.C., Sacktor, T.C. & Fenton, A.A. (2008) PKMζ maintains spatial, instrumental, and classically conditioned long‐term memories. PLoS Biol, 6, 2698‐2706.

Srinivas, S., Watanabe, T., Lin, C.S., William, C.M., Tanabe, Y., Jessell, T.M. & Costantini, F. (2001) Cre reporter strains produced by targeted insertion of EYFP and ECFP into the ROSA26 locus. BMC Dev Biol, 1, 4.

Sun, X., Bernstein, M.J., Meng, M., Rao, S., Sorensen, A.T., Yao, L., Zhang, X., Anikeeva, P.O. & Lin, Y. (2020) Functionally Distinct Neuronal Ensembles within the Memory Engram. Cell, 181, 410‐423 e417.

Takacs, V.T., Klausberger, T., Somogyi, P., Freund, T.F. & Gulyas, A.I. (2012) Extrinsic and local glutamatergic inputs of the rat hippocampal CA1 area differentially innervate pyramidal cells and interneurons. Hippocampus, 22, 1379‐1391.

Tsokas, P., Grace, E.A., Chan, P., Ma, T., Sealfon, S.C., Iyengar, R., Landau, E.M. & Blitzer, R.D. (2005) Local protein synthesis mediates a rapid increase in dendritic elongation factor 1A after induction of late long‐term potentiation. J Neurosci, 25, 5833‐5843.

Tsokas, P., Hsieh, C., Yao, Y., Lesburgueres, E., Wallace, E.J., Tcherepanov, A., Jothianandan, D., Hartley, B.R., Pan, L., Rivard, B., Farese, R.V., Sajan, M.P., Bergold, P.J., Hernandez, A.I., Cottrell, J.E., Shouval, H.Z., Fenton, A.A. & Sacktor, T.C. (2016) Compensation for PKMzeta in long‐term potentiation and spatial long‐term memory in mutant mice. eLife,

Tsokas, P., Ma, T., Iyengar, R., Landau, E.M. & Blitzer, R.D. (2007) Mitogen‐activated protein kinase upregulates the dendritic translation machinery in long‐term potentiation by controlling the mammalian target of rapamycin pathway. J Neurosci, 27, 5885‐5894.

Tsokas, P., Rivard, B., Hsieh, C., Cottrell, J.E., Fenton, A.A. & Sacktor, T.C. (2019) Antisense Oligodeoxynucleotide Perfusion Blocks Gene Expression of Synaptic Plasticity‐related Proteins without Inducing Compensation in Hippocampal Slices. Bio Protoc, 9.

Vazdarjanova, A., Ramirez‐Amaya, V., Insel, N., Plummer, T.K., Rosi, S., Chowdhury, S., Mikhael, D., Worley, P.F., Guzowski, J.F. & Barnes, C.A. (2006) Spatial exploration induces ARC, a plasticity‐related immediate‐early gene, only in calcium/calmodulin‐dependent protein kinase II‐positive principal excitatory and inhibitory neurons of the rat forebrain. The Journal of comparative neurology, 498, 317‐329.

Vogt‐Eisele, A., Kruger, C., Duning, K., Weber, D., Spoelgen, R., Pitzer, C., Plaas, C., Eisenhardt, G., Meyer, A., Vogt, G., Krieger, M., Handwerker, E., Wennmann, D.O., Weide, T., Skryabin, B.V., Klugmann, M., Pavenstadt, H., Huentelmann, M.J., Kremerskothen, J. & Schneider, A. (2013) KIBRA (KIdney/BRAin protein) regulates learning and memory and stabilizes Protein kinase Mzeta. Journal of neurochemistry.

Volk, L.J., Bachman, J.L., Johnson, R., Yu, Y. & Huganir, R.L. (2013) PKM‐zeta is not required for hippocampal synaptic plasticity, learning and memory. Nature, 493, 420‐423.

Wang, S., Sheng, T., Ren, S., Tian, T. & Lu, W. (2016) Distinct Roles of PKCiota/lambda and PKMzeta in the Initiation and Maintenance of Hippocampal Long‐Term Potentiation and Memory. Cell Rep, 16, 1954‐1961.

Wesierska, M., Adamska, I. & Malinowska, M. (2009) Retrosplenial cortex lesion affected segregation of spatial information in place avoidance task in the rat. Neurobiology of learning and memory, 91, 41‐49.

Westmark, P., Cj, W., Wang, S., Levenson, J., Kj, O.R., Burger, C. & Malter, J. (2010) Pin1 and PKMζ sequentially control dendritic protein synthesis. Science Signaling.

