## Supplementary figures for "Persistent increases of PKMζ in memory-activated neurons trace LTP maintenance during spatial long-term memory storage"

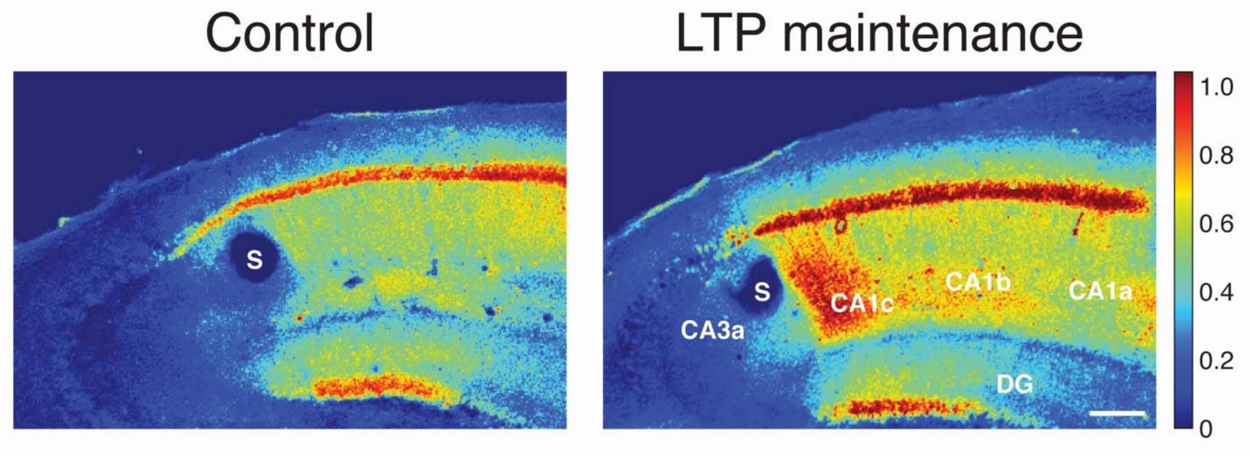


Supplementary Fig. 1. Low magnification images of non-tetanized (left) and tetanized (right) hippocampal slices show increases of PKMζ-immunointensity localized to sites of projections of Schaffer collaterals stimulated in CA3a radiatum. Increases are largest in CA1c and diminish in CA1b and CA1a. No increases are observed in regions outside the projections, including CA3a and DG. Scale bar = 100 µm; s, placement of stimulating electrode.

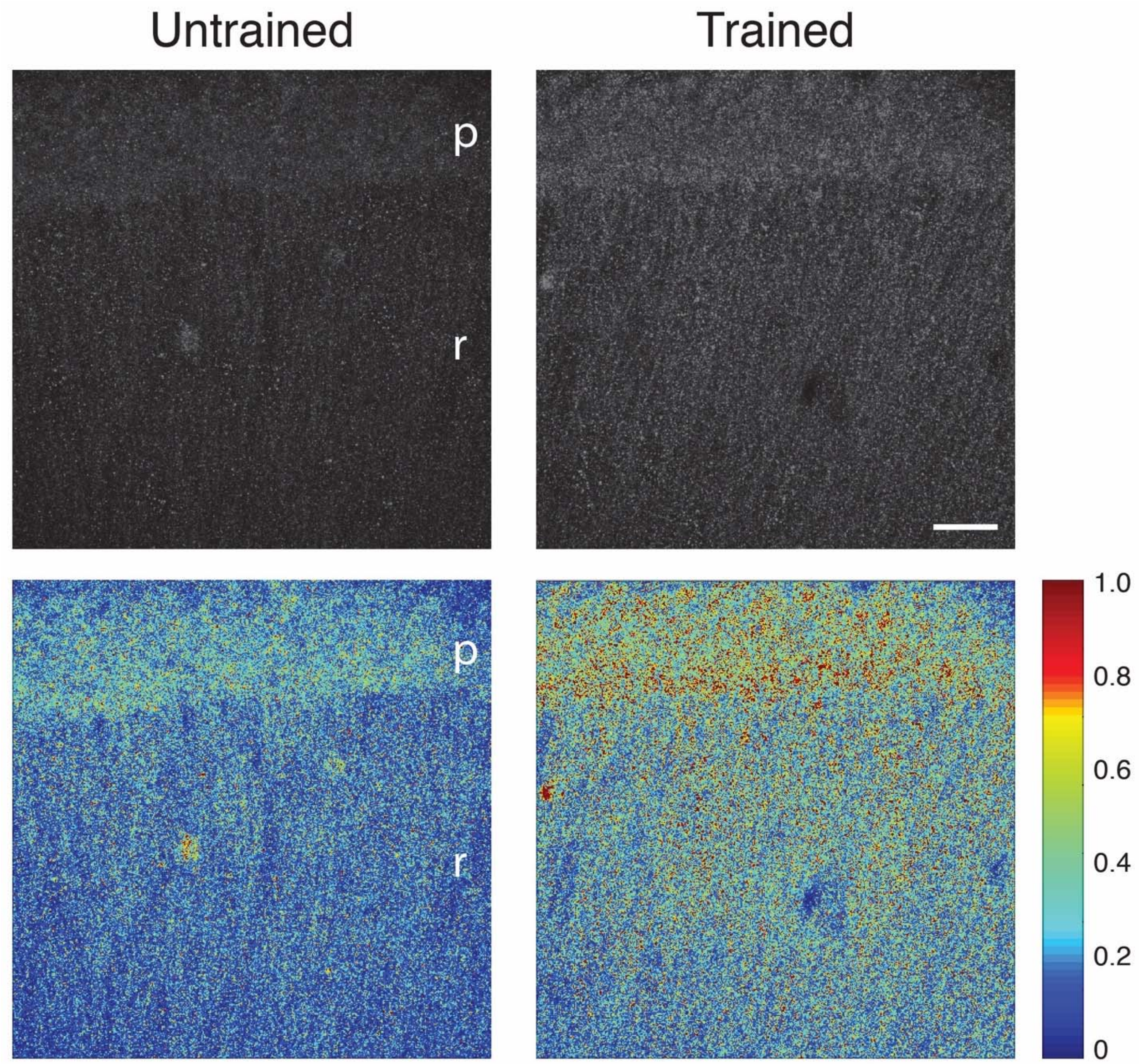


Supplementary Fig. 2. PKMζ-immunostaining in neuropil of CA1. PKMζ-immunohistochemistry of hippocampi from untrained and trained mice shows training increases PKMζ-immunostaining in p, *stratum pyramidale* and r, *stratum radiatum*. Above, gray-scaled; below, color-scaled, with colormap shown as insert at right. Scale bar = 30 μm.

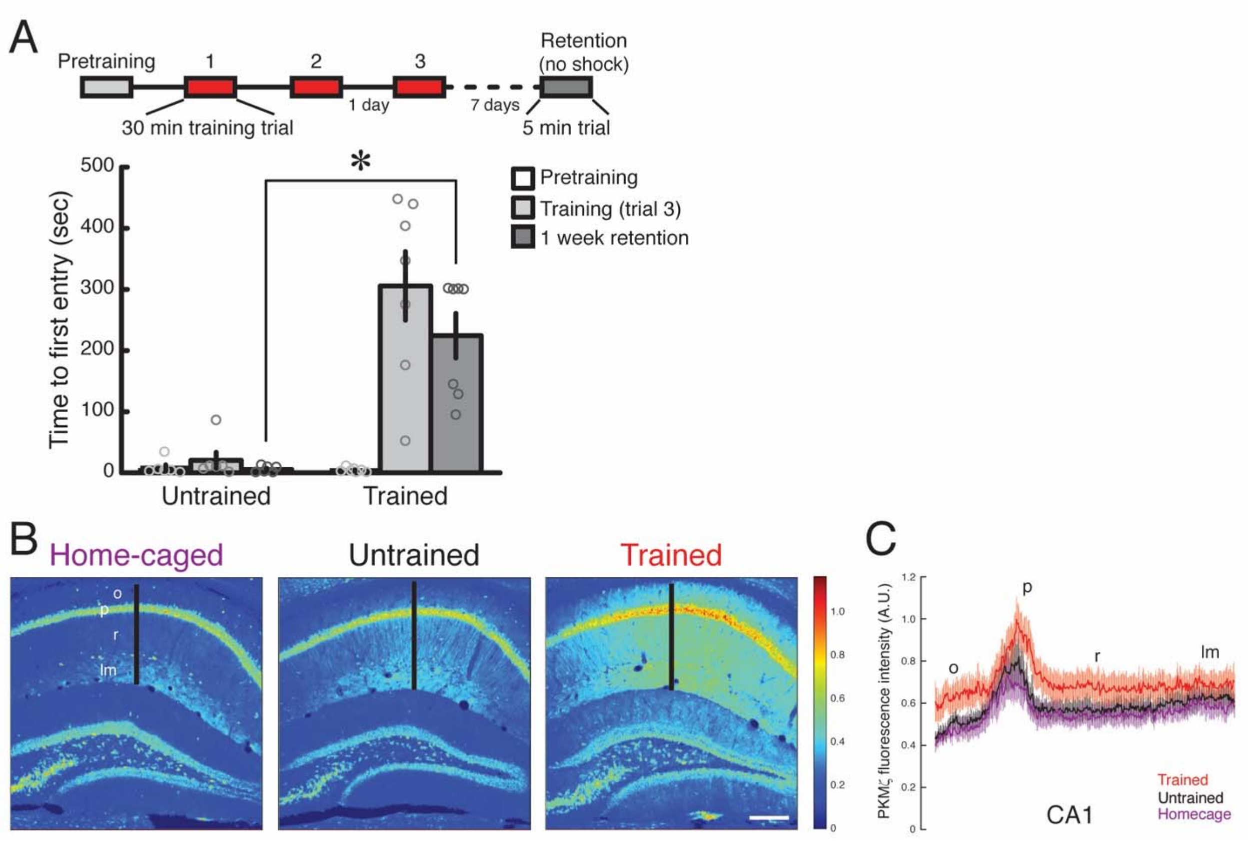


Supplementary Fig. 3. Persistent increased PKMζ in 1-week spatial long-term memory storage compared to untrained and home-caged controls. (A) Above, schematic of spatial conditioning protocol; 3 conditioning sessions separated by 1 day produce long-term memory lasting 7 days. Below, time to first entry measure of active place avoidance memory (mean ± SEM of the data shown in circles; *, statistical significance, as reported in Results). (B) Representative immunohistochemistry showing PKMζ expression in CA1 increases 1 week post-training. Black lines correspond to locations of profiles shown in (C). Colormap is shown as insert at right. Scale bar = 200 µm. (C) Profiles shows increase in PKMζ 1 week after training (red), compared to untrained (black) and home-caged control groups (purple). S*trata*: o, *oriens*; p, *pyramidale*; r, *radiatum*; lm, *lacunosum-moleculare*. Mean data ± SEM; statistical significance reported in Results.

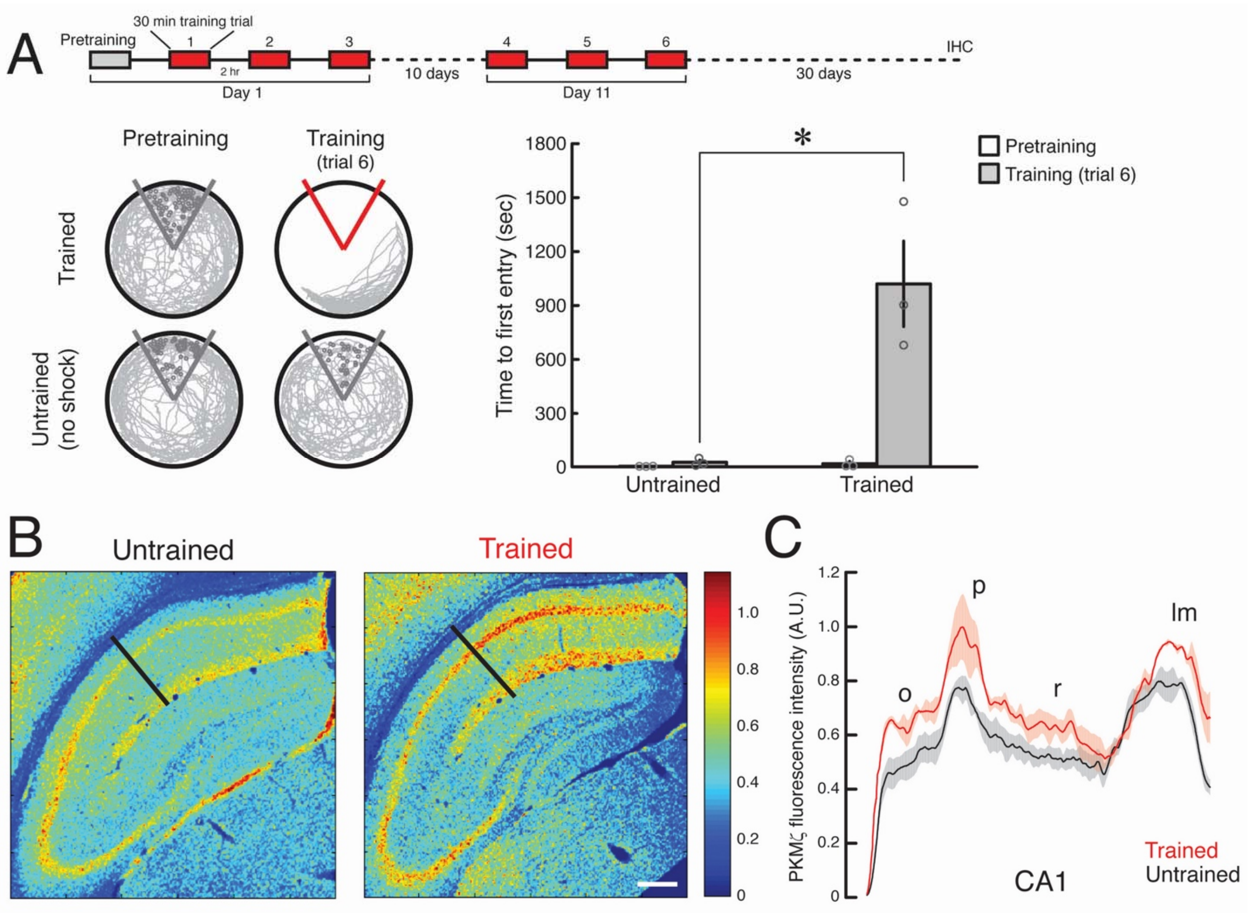


Supplementary Fig. 4. Persistent increased PKMζ during 1-month spatial memory storage without memory retention testing. (A) Above, schematic of spatial conditioning protocol with repeated conditioning separated by 10 days. Thirty days after the last training session the mice are sacrificed and analyzed for IHC without retention testing. Below left, representative 10-min periods of paths at initiation of pretraining and at the end of training. The shock zone is shown in red with shock on and gray with shock off. Gray circles denote where shocks would have been received if the shock were on. Right, time to first entry measure of active place avoidance shows conditioning at the end of training (mean ± SEM of the data shown in circles; *, statistical significance, as reported in Results). (B) Representative immunohistochemical data showing PKMζ increases in CA1 30 days post-training without memory retention testing. Black lines correspond to locations of profiles shown in (C). Colormap is shown as insert at right. Scale bar = 250 µm. (C) Profiles show increase in PKMζ 30 days after training (red), compared to untrained controls (black). S*trata*: o, *oriens*; p, *pyramidale*; r, *radiatum*; lm, *lacunosum-moleculare*. Mean data ± SEM; statistical significance reported in Results.

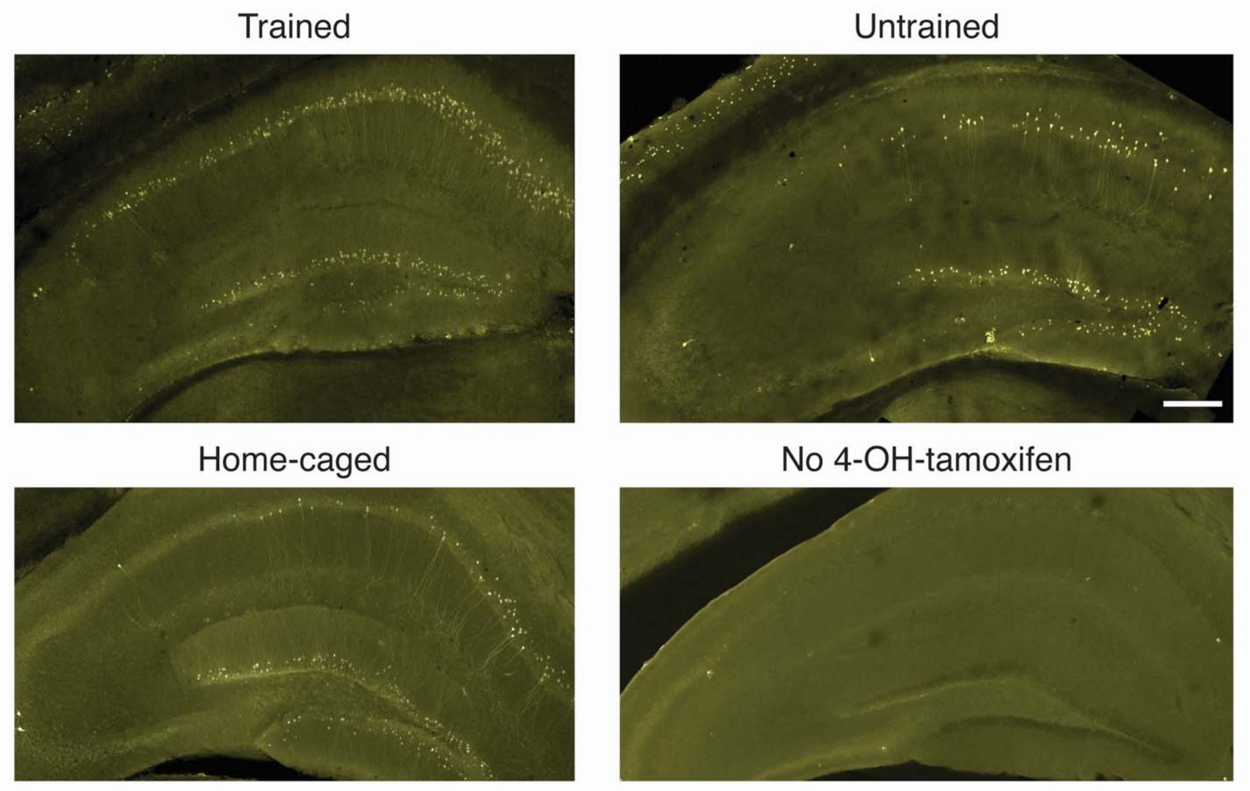


Supplementary Fig. 5. Representative image of eYFP+ cells in hippocampus of 4-OH-tamoxifen-injected trained mice in experiments described in Fig. 6, compared to eYFP+ cells in 4-OH-tamoxifen-injected untrained mice, 4-OH-tamoxifen-injected home-caged mice, as well as home-caged mice not injected with 4-OH-tamoxifen. Untrained mice were placed in the apparatus for the equivalent period of time as trained mice, but not shocked, and were injected with 4-OH-tamoxifen at the equivalent time. In all conditions, CA1 expresses more memory-tagged eYFP+ cells than DG, which is more than CA3. No 4-OH-tamoxifen shows minimal eYFP expression. The percent ratios of eYFP+ cells to total CA1 pyramidal cell bodies as defined by central DAPI staining (eYFP+ cells/total cells X 100; mean ± SEM) are as follows. CA1: trained, 11.3 ± 0.6; untrained, 4.2 ± 0.3; home-caged, 2.8 ± 0.2. DG: trained, 3.8 ± 0.1; untrained, 2.6 ± 0.2; home-caged, 3.0 ± 0.07 (n's = 5 for all groups). The 2-way ANOVA for CA1 and DG with repeated measurement reveals main effects of treatment (trained, untrained, and home-caged; $F_{2,12} = 102.8$, $P < 0.00001$), region (CA1 and DG, $F_{1,12} = 234.5$, $P < 0.00001$), and their interaction ($F_{2,12} = 143.8$, $P < 0.00001$). Post-hoc tests show that in CA1, the ratio of eYFP+ cells to total pyramidal cells of the trained group is higher than either untrained or home-caged groups, and the untrained group is higher than the home-caged group ($P < 0.00001$, $P < 0.00001$, and $P = 0.006$, respectively). In DG, the ratio in the trained group is higher than in the untrained control ($P = 0.01$), but there is no difference between trained and home-caged groups ($P = 0.06$), or between untrained and home-caged groups ($P = 0.42$). In addition, the percent ratio in the trained group in CA1 is higher than in the trained group in DG ($P < 0.00001$). Scale bar = 250 µm.
